# CRISPR/Cas9-mutagenesis reveals that varying dependence on HSF1 is associated with differences in coral heat tolerance

**DOI:** 10.64898/2026.04.01.714264

**Authors:** Natalie Swinhoe, Amanda I Tinoco, Dania Nanes Sarfati, Catherine F. Henderson, Griffin P. Kowalewski, Emily K. Meier, John M. Urban, Shumpei Maruyama, Evan C. Lawrence, Ryan E. Hulett, Ty R. Engelke, Jamie Craggs, Line K. Bay, Phillip A. Cleves

## Abstract

Coral reefs face declines due to increasing water temperatures associated with climate change. Major research efforts have focused on determining the mechanisms corals can use to adapt to heat stress and identifying molecular indicators for this adaptation. CRISPR/Cas9-based genomic editing promises a new avenue to study gene function in corals; however, these methods are limited by the annual spawning of corals in the wild. Here, we shifted spawning of the reef-building coral *Galaxea fascicularis* to access gametes multiple times a year in the lab. We discovered the remarkable plasticity and programmability in coral spawning, which enabled the development of a genetically tractable model coral. To investigate the molecular responses of corals to heat stress, we profiled transcriptional changes in heat-tolerant *G. fascicularis* and heat-sensitive *Acropora millepora* during acute heat stress. Comparison of the transcriptional responses to heat stress in larvae of the two species revealed that *A. millepora* has a stronger magnitude of the early heat stress response than *G. fascicularis*. This increased response in *A. millepora* included the upregulation of the conserved transcriptional regulator of heat stress response, *Heat Shock Transcription Factor 1* (*HSF1*), and its predicted targets. CRISPR/Cas9 mutagenesis of HSF1 in both species showed that the heat-tolerant *G. fascicularis* is less dependent on HSF1 than *A. millepora* for survival during acute heat stress. These results suggest that differences in *HSF1* expression after heat exposure contribute to variation in coral heat tolerance and may be used as biomarkers to predict heat tolerance in wild corals.

## Introduction

Reef-building corals create ecosystems of economic and environmental value owing to a nutritional endosymbiosis with dinoflagellate algae (family Symbiodiniaceae)^1^. Over the last four decades, increasing global ocean temperatures have led to the deterioration of coral reefs worldwide^2–5^. Heat stress can cause the symbiotic relationship between algal symbiont and host to break down, a process known as “coral bleaching”. If the heat exposure is extreme or prolonged and the symbiosis cannot be reestablished, corals eventually die. However, severe heat waves can also cause corals to die without first bleaching, a distinct phenotype from bleaching-induced mortality^6^. The anticipated increase in frequency and severity of marine heat waves due to climate change^7,8^ indicates that direct mortality without bleaching is an accelerating threat to corals. As such, there is a need to understand the molecular basis of coral heat tolerance and its variation to predict the future of coral adaptation to climate change and to develop molecular methods to identify and track thermally tolerant coral species.

Understanding the genetic bases for variation in heat and bleaching tolerance requires a foundational molecular understanding of these stress responses in corals (see reviews in refs. 9,10). Experimental and observational studies have shown that variation in coral heat and bleaching tolerance is associated with both environmental and genetic factors^11–17^. Among the associated genetic factors are genomic loci with putative roles in the general heat stress response, including the unfolded protein response (UPR) (e.g., *Hsp70*, *Hsp90*, and the heat shock co-chaperone *sacsin*)^10,12,18–20^. Additionally, gene-expression studies also implicate an early transcriptional upregulation of a large suite of genes during short-term heat stress in corals and other cnidarians^21–25^. Interestingly, variation in the magnitude of upregulation of this early heat stress response has been correlated with variation in heat tolerance within and between coral species^21,26^.

A key regulator of the early heat response in eukaryotes is the transcription factor, *Heat Shock Factor 1* (*HSF1*). HSF1 is transcriptionally upregulated within a few hours of heat onset in cnidarians, and CRISPR/Cas9 mutagenesis has revealed that HSF1 is required for heat survival in *Acropora millepora* larvae^22,27^. Acroporid corals are generally the most heat sensitive corals on the reef and are among the first to die during heat stress^15^, while corals in the genus *Galaxea* are known to be more resistant to heat-induced bleaching in the wild^15,28–30^. The contrast in heat tolerance between these two taxa offers an experimental opportunity to compare the genetic pathways regulating heat tolerance.

Using CRISPR/Cas9 genome editing, it should be possible to identify the molecular basis of variation in heat tolerance through the genetic manipulation of candidate genes and pathways^31^. However, microinjection of CRISPR/Cas9 reagents into single-cell zygotes is restricted by access to coral gametes and relies on natural spawning events, which typically occur only once or twice a year^27,27,32–34^. Therefore, progress towards functional characterization of genes in corals has been limited. Recently, the ability to spawn corals in laboratory culture has become possible for some species using closed artificial mesocosms, where gametogenesis can be induced by following natural environmental parameters (e.g., temperature cycles, solar and lunar irradiance levels, sunset and sunrise times)^35–37^. Recent research indicates that it is possible to shift coral spawning by 6 months and retrieve viable offspring^38^. Combining CRISPR/Cas9 methods with year-round and predictable access to coral gametes in the laboratory would greatly accelerate the pace with which reverse genetics can elucidate gene function in corals.

Here, we developed the heat-tolerant *Galaxea fascicularis* as a genetically tractable reef-building coral. We chose this species as it has a wide distribution throughout the Indo-Pacific and Red Sea and it can be easily grown in aquaria^39,40^. We shifted the gametogenic cycles of colonies in separate aquaria to induce predictable and reliable spawning at staggered intervals during the year in the laboratory. To investigate the molecular regulation of coral heat tolerance, we generated a new megabase-scale genomic assembly for *G. fascicularis* and performed RNA sequencing of both *A. millepora* and *G. fascicularis* larvae during heat stress. We identified a large difference in the magnitude of the early heat stress transcriptional response between *G. fascicularis* and *A. millepora*, which correlates with the reported differences in heat tolerance of these species in the wild. This response included *HSF1* and predicted targets. We developed microinjection methods to produce mRNA-based gene overexpression and CRISPR/Cas9-based mutagenesis in *G. fascicularis*. Using these new genetic methods, we found that *G. fascicularis* larvae are less dependent on *HSF1* for survival to heat stress than *A. millepora* larvae. Our results suggest that the HSF1 pathway underlies variation in heat tolerance and may be an avenue for heat adaptation in corals.

## Results

### Year-round spawning of *G. fascicularis* in the laboratory

To make *G. fascicularis* a genetically tractable model system, we established year-round spawning in the lab and developed methods for gene editing and rearing modified animals for experimentation (Figure 1A). Four cohorts of *G. fascicularis* colonies were sourced from the Great Barrier Reef (GBR) in October 2021, February 2022, September 2022, and November 2023 (Table S1), and were housed in four premanufactured aquaria (‘spawning systems’) with programmable temperature and light controls^35^. Initially, each aquarium was set to match the solar and lunar photoperiods, lunar intensity, and average temperature observed in Moore Reef, GBR in 2021 (see Materials and Methods) (Figure 1B). These conditions have been shown to induce *G. fascicularis* spawning over a 12-month cycle on days six, seven, and eight post full moon during the predicted spawning month^37^. The first *G. fascicularis* cohort received in October 2021 spawned in the laboratory in November 2021, in synchrony with predicted spawning of wild *G. fascicularis* colonies from the GBR (Table S2).

**Figure 1.**
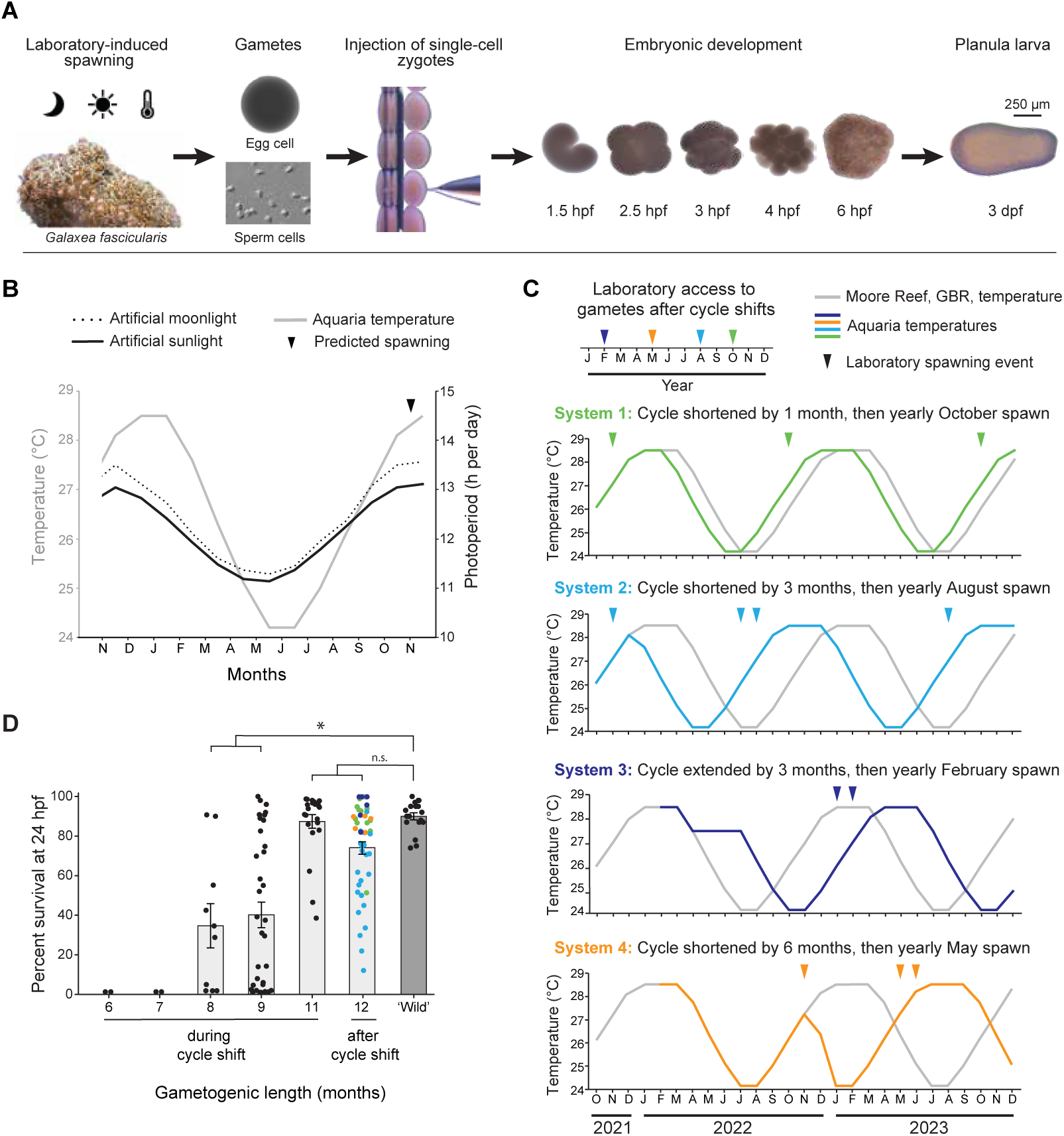
Year-round access to *G. fascicularis* gametes is created by shifting spawning cycles. **(A)** Strategy for the development of *G. fascicularis* as a tractable system, including laboratory access to coral gametes, microinjection of single-cell zygotes, and rearing of mutant larvae for experimentation. **(B)** Modified seasonal temperature, solar photoperiod, and lunar photoperiod parameters from Moore Reef, GBR, that were shifted to induce coral spawning at predictable intervals (black arrowhead). **(C)** Strategy for predictable access to coral gametes four times a year by shifting environmental parameters in four aquarium systems. Only temperature is shown for each system, although sun, moon, and photoperiod cycles were shifted synchronously. Gametogenic length is the distance between spawns (arrows), except in cases of consecutive monthly spawnings. The unchanged temperature profile of Moore Reef, GBR is shown in gray, as in (B). **(D)** Percent survival of animals 24 hours post-fertilization (hpf) using gametes spawned after different gametogenic cycle lengths. Individual fertilizations are shown (dots) with mean and standard error. Colors correspond to the systems in (C). ‘Wild’ fertilizations used gametes from coral colonies collected from the GBR in October 2021, held in the aquarium systems for one month, and spawned in November 2021. Statistical significance was determined by a Kruskal-Wallis test followed by a Dunn’s multiple comparison test with Bonferroni correction. (* = p ≤ 0.05).

Next, we adjusted the environmental parameters in each of the spawning systems to induce regular laboratory spawning four times a year. The shifts were accomplished by removing one month of summer (System 1), removing three months of summer (System 2), adding three months of spring (System 3), or removing six months of spring and summer (System 4) (Figure 1C). Each of these shifts resulted in spawning during the predicted month with 79%, 74%, 32%, and 9% of corals spawning in each of the four systems, respectively (Figure 1C). Interestingly, the corals in Systems 2, 3, and 4 released gametes either the month before or the month after the predicted spawning month, resulting in a split spawn following the shift. After determining optimal conditions for fertilization of gametes (Figure S1A and 1B), we asked if shortening the length of gametogenesis impacted coral reproductive output by assessing larvae survival (Figure 1D). Fertilizations of gametes from a shortened gametogenic cycle had lower survival compared to the standard 12-month cycle (Figure 1D). Specifically, no surviving larvae were produced from the crosses from animals with a six- or seven-month cycle, and a significant reduction of surviving larvae was observed from the crosses from animals with an eight- or nine-month cycle. There was no significant difference in larvae survival from a shortened eleven-month gametogenic cycle (Figure 1D). The number of eggs per bundle was also decreased after the 6-month shift but then recovered after returning to the 12-month cycle (Figure S1C). These results indicate reproductive limits to the ability to shorten the gametogenic cycle of *G. fascicularis*. After the spawn from the shifted cycles, we reverted each system back to a 12-month gametogenic cycle. The first 12-month gametogenic cycle after the shift, 80%, 100%, 67%, and 26% of coral colonies spawned from the first spawning cohort, respectively (Table S2). Fertilization and development of *G. fascicularis* embryos from gametes produced after a 12-month gametogenic cycle proceeded normally^41,42^ (Figure S1D-S1F). These results demonstrate that it is possible to induce year-round coral spawning, increasing access to zygotes for genetic editing and other experiments.

### Reduced heat stress response in *G. fascicularis* larvae compared to *A. millepora* larvae

Coral species from the genus *Acropora* have been reported to be more heat sensitive than those in the genus *Galaxea*^15^. To investigate the transcriptional basis of these varying heat stress response phenotypes, we first improved the genomic reference of *G. fascicularis* by generating a mega-base scale genome (Tables S3-S6). Next, we performed an RNA-sequencing time course during heat stress at 34 °C of both *A. millepora* and *G. fascicularis* larvae starting at 3 days post fertilization (dpf). To compare the early heat stress responses between the two species, we identified genes that were significantly and highly differentially expressed (*p*-adj ≤ 0.05, log_2_ fold-change ≥ 2 or ≤ −2) at 1.5, 3, 6, 12, and 24 h of heat compared to 0 h within each species (Dataset S1, Dataset S2). Strikingly, we found that *G. fascicularis* had fewer differentially expressed genes at all but one timepoint compared to *A. millepora* (Figure 2A). To explore these differences further, we identified the top 100 genes with the largest upregulation at 3 h post heat stress in each time course, normalized each gene to its respective 0 h timepoint, and sorted each gene set according to the fold change from 0 to 3 h (Figure 2B and 2C). These genes show a dramatic reduction in the magnitude of upregulation of expression at 3 h in *G. fascicularis* compared to *A. millepora*. To ask if the same genes were deployed during the heat-stress response between each species, we identified 14,431 one-to-one orthologs between *A. millepora* and *G. fascicularis* using reciprocal best BLAST. We compared the fold changes for the one-to-one orthologs that were upregulated at 3 h in *A. millepora* between the two species. Interestingly, the majority (167 of 187) of upregulated orthologs at 3 h in *A. millepora* were not significantly upregulated at 3 h in *G. fascicularis* (Dataset S3). Gene frontloading, where gene expression is higher at baseline conditions, has been correlated with the lack of transcriptional response during stress in corals^21,26^. To test if frontloading could explain the lack of transcriptional upregulation of these orthologs in *G. fascicularis*, we plotted the normalized expression of the one-to-one orthologs at 0 h and 3 h in both species (Figure 2D). We found no evidence of frontloading of these genes by mRNA in *G. fascicularis* compared to *A millepora* at 0 h, suggesting that global frontloading of stress response genes cannot explain the differences across these two species.

**Figure 2.**
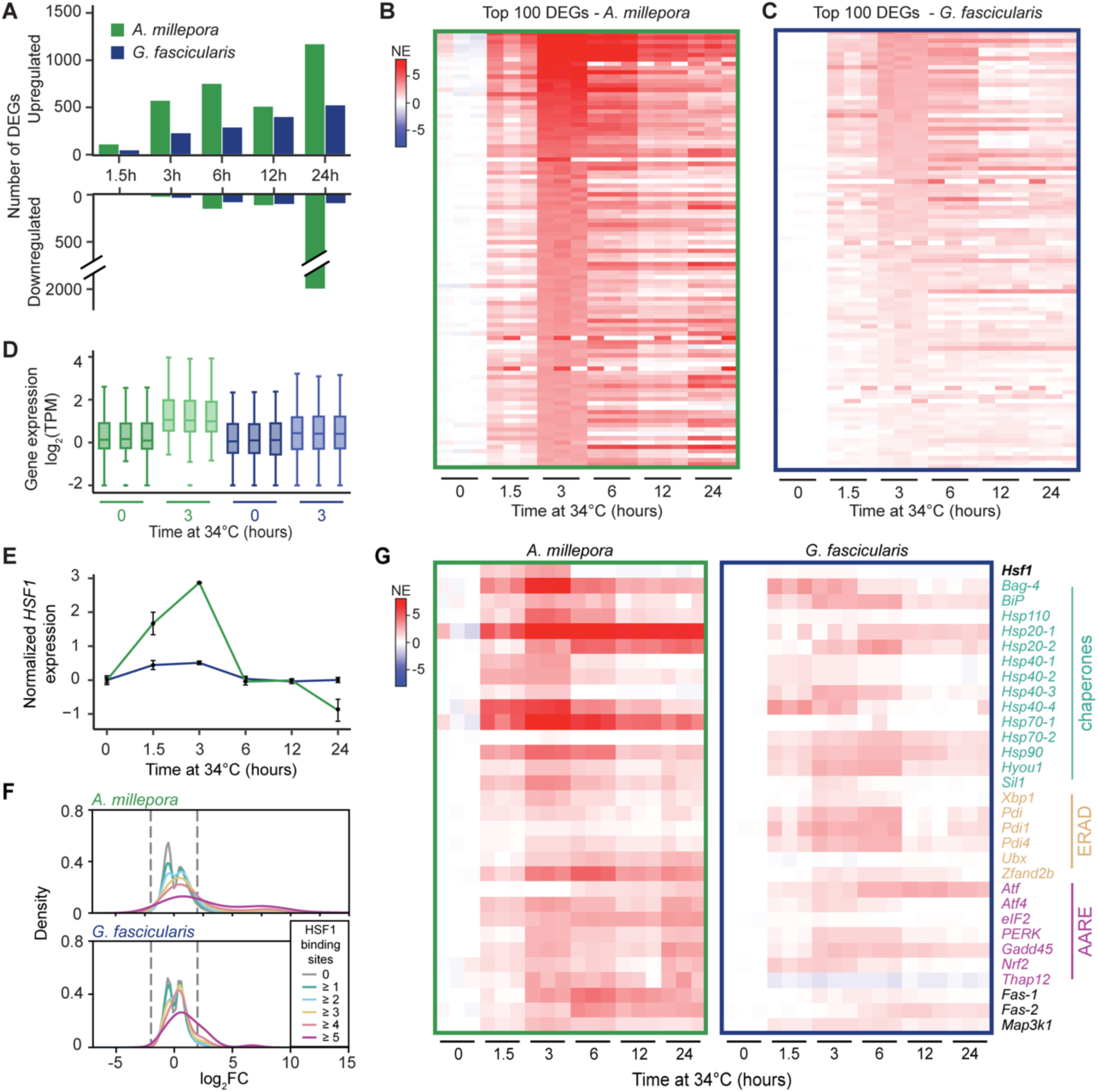
Differential upregulation of HSF1 and UPR pathways during heat stress across species. **(A)** Total DEGs (*p*-adj ≤ 0.05, log_2_ fold-change ≥ 2 or ≤ −2) identified at each time point compared to 0 h per species. **(B-C)** Top 100 differentially upregulated genes at 3 h of heat exposure in *A. millepora* (**B**) or *G. fascicularis* (**C**). Color bar corresponds to log_2_ expression of each of three replicates per time point normalized by the mean expression at time point 0 h for each species. The increased intensity of the response observed in *A. millepora* is recapitulated at all time points. NE, normalized expression. **(D)** Normalized expression (log_2_(TPM)) of upregulated genes in *A. millepora* at 3 h of heat exposure that have an ortholog in *G. fascicularis* show no overall frontloading in *G. fascicularis* larvae. **(E)** Normalized expression of *HSF1* in both species, showing significant upregulation at 1.5 and 3 h in *A. millepora* but not in *G. fascicularis*. **(F)** Smoothed density histograms for genes with different numbers of HSF1 binding sites as a function of log_2_ fold change expression (log_2_FC) showing a positive relationship between higher number of binding sites and higher upregulation in response to 3h of heat stress in *A. millepora* (top) and *G. fascicularis* (bottom). Dashed lines show significance cut off for log_2_FC. **(G)** Heatmaps of selected orthologous genes involved in the unfolded protein response in *A. millepora* (left) and *G. fascicularis* (right). ERAD, ER-associated degradation; AARE, amino acid response element; NE, normalized expression. For all panels, green: *A. millepora,* blue: *G. fascicularis*.

Our previous research found that the transcription factor, HSF1, is required for survival during heat stress in *A. millepora* larvae^27^. We confirmed that *HSF1* is indeed significantly upregulated in *A. millepora* at 3 h (Figure 2E). Strikingly, *HSF1* was not significantly upregulated during heat stress at any time point in *G. fascicularis*. To determine if predicted *HSF1* targets are also dampened in *G. fascicularis* compared to *A. millepora,* we identified genes in both species that have predicted HSF1 binding sites in their putative promoter regions defined as the 500 bp immediately upstream of their transcription start sites^22^. Then, we compared the number of HSF1 sites and the degree of upregulation after 3 h of heat exposure in each species. Genes with more HSF1 sites were more likely to be upregulated during heat stress in both species, and the magnitude of upregulation of these putative HSF1-target genes was higher in *A. millepora* than *G. fascicularis* (Figure 2F). Together, these results suggest that *HSF1* and some of its downstream targets are upregulated in *A. millepora* during heat stress, but this response is dampened in *G. fascicularis*.

Because the HSF1 pathway is a key pathway in the unfolded protein response, we hypothesized that other pathways of the UPR might also be differentially upregulated between these two coral species. We searched the one-to-one ortholog list for other members of the UPR that were upregulated in either species at any heat stress timepoint. We found several chaperones, including various Heat Shock Proteins and other core members of the UPR, including *XBP1*, *PERK*, *ATF4*, and *NRF2*^43–46^. Strikingly, many of these genes were upregulated in *A. millepora* and *G. fascicularis*, but their degree of upregulation was less in *G. fascicularis* (Figure 2G, Dataset S1, Dataset S2). These results show that *A. millepora* larvae have a much stronger UPR during heat stress compared to *G. fascicularis* larvae.

### CRISPR/Cas9 mutagenesis reveals differences in dependence on *HSF1* during heat stress between coral species

The differences in magnitude of the early heat stress response and *HSF1* activity suggest that *A. millepora* is more reliant on *HSF1* to survive heat stress compared to *G. fascicularis* (Figures 2E-2G). To test this hypothesis, we developed methods for CRISPR/Cas9 mutagenesis in *G. fascicularis*. Previously, we found that co-injection of CRISPR/Cas9 reagents with a fluorescent injection indicator (Alexa Fluor 488-dextran) into 1-cell zygotes of *A. millepora* allowed for efficient sorting of highly mutant individuals at 24 hours post fertilization (hpf) using green fluorescence^27,34^. However, *G. fascicularis* eggs have endogenous green fluorescence (Figure S2A), making it difficult to distinguish autofluorescence from the 488-dextran injection indicator at 24 hpf. To overcome this challenge, we injected newly fertilized zygotes with capped mRNA encoding a cyan fluorescent protein (CFP), mTurquoise2 (mTQ2)^47^ (Figures S2B and S2C), which is scorable at 24 hpf as *G. fascicularis* larvae do not have endogenous cyan fluorescence (Figure S2A). There was no significant difference in survival at 24 h between animals injected with phenol red or the combination of phenol red and mTQ2 mRNA, indicating that injecting capped mRNA did not impair larval development or survival (Figure S2D and Table S7). Of the surviving larvae injected with mTQ2 mRNA, between 89% and 95% were scored as positively injected by the presence of cyan fluorescence (Figures S2C, S2E, and Table S7). To determine the duration of overexpression, a subset of larvae with uniform cyan fluorescence at 24 hpf were imaged live intermittently until 23 dpf. Cyan fluorescence was detectable but diminished by 23 dpf (Figures S2F and S2G). These results indicate that mRNA can be used to transiently overexpress target genes in *G. fascicularis*.

Next, we used mTQ2 mRNA as an injection indicator along with CRISPR/Cas9 reagents to generate mutations in *HSF1* in *G. fascicularis* (Figure S2H). First, we designed two sgRNA guides targeting exon 3 and exon 4 of *HSF1* (Figure 3A and Table S8). Over three days of spawning, we injected three experimental replicates of 1-cell zygotes with a mixture of HSF1 sgRNA/Cas9 complexes, 488-dextran, and mTQ2 mRNA. Surviving larvae were sorted as successfully injected based on the presence of cyan fluorescence at 24 hpf (Figure S3 and Table S9) and reared at 27 °C until 3 dpf. We then quantified the mutation frequencies of successfully injected larvae at 3 dpf by Sanger sequencing PCR products spanning each sgRNA target site (Table S10). Most of the larvae sequenced (34 of 39) were completely mutant at each sgRNA site (Figure 3B and Table S10). Approximately 75% of the larvae (30 of 39) had at least one large deletion (∼400 bp) spanning from one sgRNA site to the other, and 41% (16 of 39) had at least 50% large deletions (Table S10), showing that it is possible to make large genomic deletions by using two closely spaced sgRNAs. Using this technique, we were able to generate *G. fascicularis* animals that were highly mutant for *HSF1* at frequencies comparable to previously published work in *A. millepora* (Figure 3B)^27^.

**Figure 3.**
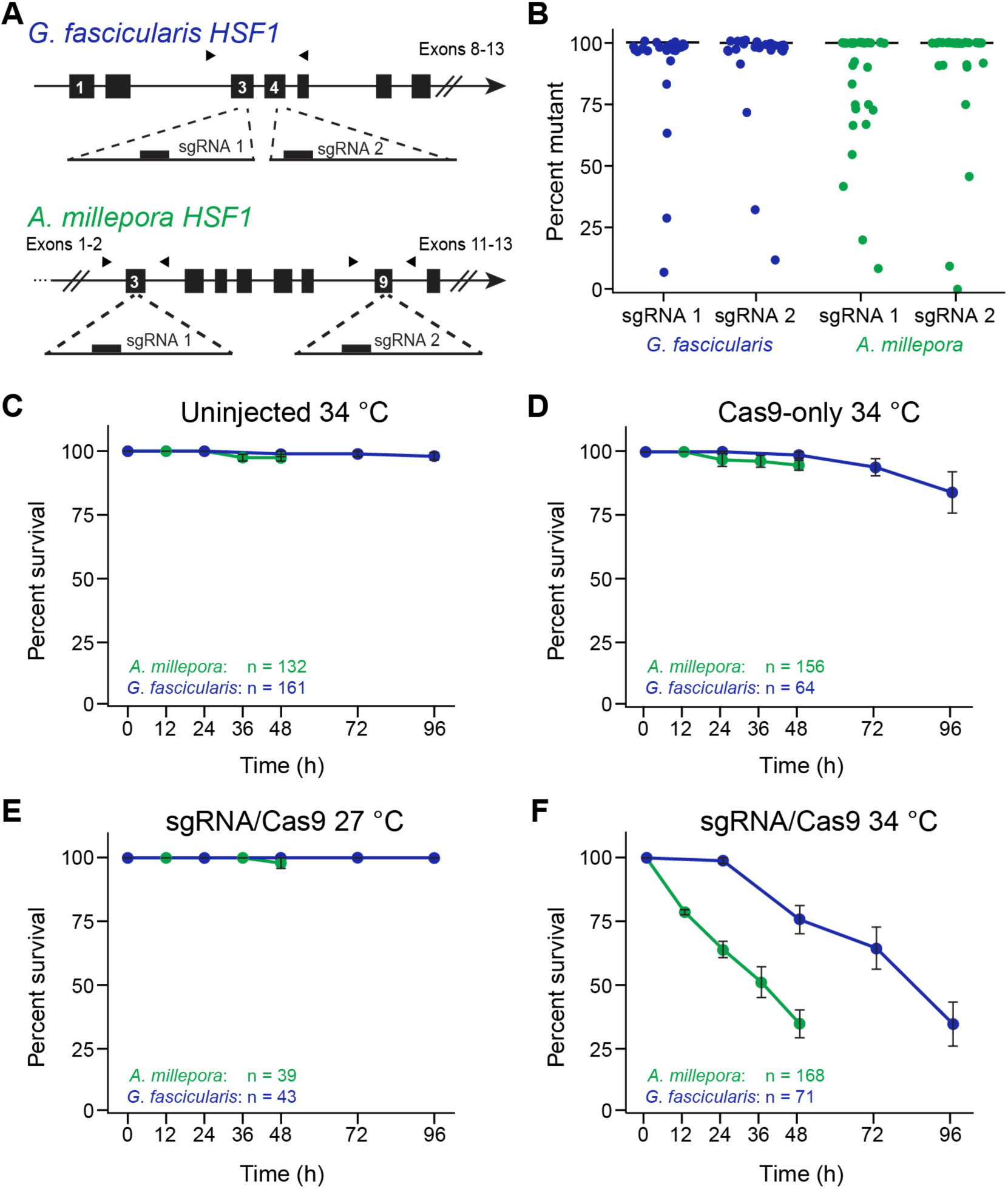
CRISPR/Cas9 mutagenesis reveals species-specific differences in *HSF1* dependence during heat stress. **(A)** Schematic of *HSF1* in *G. fascicularis* and *A. millepora* with the locations of sgRNA sites. Exons 1-7 and 3-9 (black boxes) are shown for *G. fascicularis* and *A. millepora*, respectively, as are the primers used for Sanger sequencing (black arrowheads). **(B)** Mutation percentages at each sgRNA site of *G. fascicularis* (blue) and *A. millepora* (green) larvae injected at the 1-cell stage with sgRNA/Cas9 complexes targeting *HSF1*. The median percent of mutation sequences for each species at both sgRNA sites is shown (black bar). **(C-D)** Percent survival of larvae that were either uninjected **(C)** or injected at the 1-cell stage with Cas9 only **(D)** exposed to 34 °C for 96 h. **(E-F)** Percent survival of larvae injected at the 1-cell stage with sgRNA/Cas9 complexes targeting *HSF1* at 27 °C **(E)** or 34 °C **(F)** for 96 h. For **(C-F)**, mean and standard errors for three independent injection replicates from three spawning days are shown as are the total number of larvae across all three replicates. Experiments began when the larvae were 3 dpf. For **(B-F)**, only larvae scored as successfully injected at 24 hpf were used. Data for *A. millepora* experiments are from ref. 27.

To determine if *HSF1* is required for *G. fascicularis* survival in heat stress, we placed uninjected, Cas9-only injected, and sgRNA/Cas9-injected larvae at 34 °C for 96 h, in the same manner as was previously described for *A. millepora*^27^. There was minimal mortality of uninjected or Cas9-only injected controls at 34 °C (Figure 3C and 3D, blue), as well as no mortality at 27 °C of larvae injected with sgRNA/Cas9 targeting *HSF1* (Figure 3E, blue). However, the larvae injected with sgRNA/Cas9 targeting *HSF1* began to die after 48 h at 34 °C (Figure 3F, blue). By 96 h, only 35% of larvae injected with sgRNA/Cas9 targeting *HSF1* survived, compared to 100% and 84% of uninjected and Cas9-only controls, respectively. These results show that *HSF1* is required for survival during heat stress for *G. fascicularis* larvae.

To evaluate if *G. fascicularis* and *A. millepora* differ in their dependence on *HSF1* upon heat stress, we compared the death rates due to heat stress of *G. fascicularis* larvae with mutations in *HSF1* to previously published data of *A. millepora* from ref. 27 (Figures 3C-3F, green). Strikingly, *A. millepora* larvae with mutations in *HSF1* died rapidly upon heat stress, relative to *G. fascicularis* larvae with equivalent mutation rates in *HSF1* (Figure 3B). By 24 h of heat stress, 38% of *A. millepora* larvae had died, whereas no mortality was observed in *G. fascicularis* larvae (Figure 3F). By 48 h of heat stress, 67% of *A. millepora* larvae had died, whereas comparable mortality in *G. fascicularis* larvae was not observed until 96 h. As a control, we also injected *G. fascicularis* zygotes with and without mTQ2 mRNA in an independent experiment (Figures S4A, S4B and Table S11). We found no difference in the frequency of induced mutations or rate of death at 34 °C between the two groups, indicating that the addition of the mTQ2 mRNA was not the cause of delayed mortality in *G. fascicularis* (Figure S4C). Together, these results show that *A. millepora* larvae are more reliant on *HSF1* to survive heat stress than *G. fascicularis* larvae, indicating that variation in the HSF1 response is associated with differences in heat tolerance between coral species.

## Discussion

### Programming coral spawning: its plasticity and constraints

Here, we developed methods to induce stable and reliable spawning of *G. fascicularis* throughout the year. During these experiments, we gained insights into both the plasticity and the constraints that govern coral spawning. First, we found that *G. fascicularis* spawning time can be shifted by changes in environmental parameters (i.e., temperature and timings of sunrise, sunset, moonrise, and moonset), indicating that these cues are sufficient to reprogram *G. fascicularis* spawning. This spawning plasticity may be important in the wild across depth gradients and geographic ranges and may be a source for variation in spawning times across populations and species. Indeed, studies have identified that cryptic coral species often stratify by depth^48–50^ and that this type of speciation can occur with changes in spawning time, potentially driven by genetic variation in light-sensing G-coupled protein receptors (e.g., Rhodopsin-like proteins)^48^. However, the specific light- and temperature-sensing pathways that drive the induction of gametogenesis, synchronized spawning, and plasticity remain to be uncovered. Second, we discovered apparent limitations to the plasticity in spawning times. We found significant fecundity trade-offs when we decreased the time for gametogenesis to six months (Figure 1D). After this six-month shift, only ∼9% of the animals spawned. We hypothesize that the reduction in spawning of these animals was due to energetic limitations from an abbreviated growth phase resulting from the removal of warm months in the cycle (Figure 1C). Consistent with the hypothesis, gametogenesis for other reef-building species has been shown to start several months before spawning and after an extended growth phase ^35,37^. It may be possible to further shorten coral gametogenesis through additional food supplementation or by adjusting which days or months are removed from the spawning cycle. These types of experiments would allow a deeper understanding of the physiological constraints that govern coral reproduction. Third, although we can program coral spawning, when we extended the gametogenic cycle by 3 months to a 15-month cycle, we found that ∼60% of the animals spawned a month earlier than predicted (month 14). This result indicates that corals may have a limit to the duration that they can hold mature gametes and that asynchronous spawning can occur due to mismatches in temperature and light cues. Strikingly, the reproductive trade-offs and asynchrony caused by the shifts were rescued in the following 12-month cycle, showing that corals have remarkable plasticity in spawning timing, which has likely shaped their evolutionary trajectory^51,52^.

### *G. fascicularis* as a genetically tractable reef-building coral

The ability to control laboratory-based spawning throughout the year accelerated the development of genetic tools. Over the course of 12 spawning events in two years, we developed and implemented new CRISPR/Cas9 genome editing and mRNA-based gene overexpression methods. A key technological advance was the use of capped mRNA encoding a cyan fluorescent protein (mTQ2) as a post-injection marker to identify successfully injected animals. Using mTQ2 mRNA in conjunction with CRISPR/Cas9, we produced highly mutant *G. fascicularis* larvae. Although it was possible to inject and sort animals without the mTQ2 mRNA (Figure S4A and Table S11), the sorting is time consuming, slowing the rate of injection and sorting, making it impossible to inject enough animals for complex experiments. As many coral species have species-specific autofluorescence, the capped mRNA can be customized with other fluorescent proteins which should enable CRISPR/Cas9 in diverse coral species. Another advancement of our method was injecting two guide RNAs targeting regions that were spaced ∼500 bp apart in the genome, leading to a high frequency of genomic deletions between the two sites (Table S10). This strategy could be expanded to target specific regions of interest, such as functional domains (e.g., DNA binding domains), regulatory regions, or entire genes, opening the door for more sophisticated genomic editing experiments. Finally, as corals take several years to reproduce, the impacts of genetic modifications need to be observed in the injected generation. However, the ability to contain, spawn, settle, and rear corals in the lab will enable the ability to cross and maintain mutant and transgenic coral lines.

In addition to its use as a post-injection marker, we have shown that it is possible to overexpress proteins in corals by injecting capped mRNA. In our experiments, the mTQ2 fluorescent protein overexpression was detectable for over 20 days (Figure S2G). It remains unknown whether the prolonged duration of the cyan fluorescence was due to continued mRNA persistence or extended stability of the fluorescent protein^47^. Regardless, it should be possible to use capped mRNA to transiently overexpress target proteins at early embryonic stages and potentially during metamorphosis and settlement. This tool enables a variety of reverse genetic experiments in corals, such as overexpression of target genes^53–56^, tagging of target proteins for subcellular localization^57^, and transient knockdown experiments using RNAi^56,58–61^. The combination of laboratory spawning and these new reverse genetic tools positions *G. fascicularis* as a genetically tractable reef-building coral, setting the stage for rigorous genetic studies of diverse aspects of coral biology.

### Differential transcriptional responses to heat between *A. millepora* and *G. fascicularis*

We found an early transcriptional response after the onset of heat in both *A. millepora* and *G. fascicularis* larvae, consistent with known early heat stress responses in cnidarians^21–24,62^. Surprisingly, the number of DEGs and the magnitude of upregulation were much higher in *A. millepora* compared to *G. fascicularis* at all assessed time points, suggesting that a higher sensitivity to heat is accompanied by a greater magnitude of transcriptional response during heat stress. These differentially upregulated genes included key members of the UPR, including HSF1 and multiple putative targets (Figure 2G), indicating that the same heat stress can cause differential deployment of genes involved in maintaining protein homeostasis in different coral species. Because the magnitude of these heat stress responses correlates with the ultimate heat tolerance of the studied species, we hypothesize that it may be possible to use these genes as biomarkers for variance in heat tolerance. Given that the HSF1 response is a part of the core organismal response to heat, the negative correlation between its expression and heat tolerance may be conserved across corals. Consistent with this hypothesis, previous research has identified an association between heat tolerance and the magnitude of the stress response in adult corals of four different genera (*Favites*, *Montipora*, *Acropora*, and *Seriatopora*)^26^ and within conspecific corals that vary in heat tolerance^21^. Further research is needed to determine the extent to which these transcriptional associations hold across diverse populations, species, and genera of corals.

### Varying dependence on HSF1 between a heat-tolerant and heat-sensitive coral species

While studies have associated certain genomic loci with heat tolerance and adaptation in corals^12,18–20^, it remains unclear which genes play a functional role in heat tolerance. Here, we show that *HSF1* has a key functional association with variation in heat response in corals.

The differential reliance on HSF1 during heat stress between *A. millepora* and *G. fascicularis* could result from several molecular mechanisms. HSF1 protein levels may differ between *G. fascicularis* and *A. millepora* larvae before heat stress, which could account for the differences in the transcriptional upregulation of HSF1 and targets. Measurements of HSF1 protein levels before heat stress in both species would test this hypothesis. Similarly, unfolded proteins in *A. millepora* may accumulate faster during heat stress than in *G. fascicularis* due to the differential thermal stability of the proteome between the species, thereby requiring a quicker and more intense response of HSF1 to avoid accumulation of unfolded proteins. Direct *in vitro* measurements of each species’ proteome stability at different temperatures could test this hypothesis^63^. Finally, heat-tolerant *G. fascicularis* might have unknown HSF1-independent pathways that compensate during heat stress. Future research characterizing the differences in the HSF1 response and other heat stress programs between the species is needed to understand if such a mechanism exists.

Regardless of the precise mechanism behind the differential dependence on *HSF1* between the coral species, we identified a suite of ∼100 genes with similar transcriptional responses as *HSF1* between the two species (Dataset S4). Interestingly, many of the genes are predicted to be targets of HSF1 based on our binding site analyses, providing putative downstream effectors that may also contribute to variation in coral heat tolerance. Using the now genetically tractable coral, *G. fascicularis*, it is possible to functionally determine the roles of each of these genes in either enhancing or reducing heat tolerance. As genes controlling heat tolerance are discovered, they can be functionally tested as biomarkers to predict thermal tolerance in wild corals. Our findings set the stage for using reverse genetics to discover the genetic bases of coral heat tolerance and promise to help inform the protection of threatened coral reef ecosystems.

## Resource availability

### Lead contact

Requests for further information and resources should be directed to and will be fulfilled by the lead contact, Phillip Cleves

## Materials availability

This study did not generate new unique reagents.

## Data and code availability

All data for this paper are provided in the main text and SI Appendix. Raw sequences and metadata have been deposited in the NCBI BioProject database (accession no. XXXXXX and XXXXXX).

## Acknowledgments

We thank members of the Cleves laboratory for their helpful comments on the manuscript. We also thank members of Carnegie Science - Department of Embryology: Allison Pinder, Frederick Tan, Ted Cooper, David Ashwood, Devance Reed, Mahmud Saddiqi, Julia Baer, Harrison Curnutte, Sammasia Wilson, and Hannah Kozan. This work was funded by start-up funds from Carnegie Science (P.A.C.), an NSF-Integrative Organismal Systems (IOS) Enabling Discovery through GEnomics (EDGE) grant (2128073, P.A.C.), support from the Pew Biomedical and Marine Fellow Award (P.A.C.), support from the Australian Institute of Marine Science (L.K.B.), a grant from Revive and Restore (P.A.C.), a grant for the Moore Foundation (P.A.C.), and an International Macquarie University Research Excellence Scholarship (A.I.T.).

## Author Contributions

N.S., A.I.T., C.F.H., D.N., and P.A.C. designed the experiments. N.S., A.I.T., D.N., C.F.H, G.P.K., E.K.M., S.M., E.C.L., R.E.H., T.R.E., and P.A.C. performed the experiments. J.M.U. provided guidance on nanopore data collection and performed bioinformatic analyses for genome assembly and annotation. J.C. provided guidance on *Galaxea fascicularis* spawning. L.K.B. provided guidance on *Acropora millepora* spawning experiments. N.S., A.I.T., D.N., and P.A.C. wrote the paper. All authors reviewed and commented on the manuscript.

## Supplemental information

Document S1. Materials and Methods, Supplemental methods, Figures S1-S4, Tables S1-S12, and supplemental references.

Dataset S1. *Galaxea fascicularis* genes with log2FC values and adjusted p-values for all timepoints compared to 0 h and predicted HSF1 binding sites in the 500bp region upstream of each gene.

Dataset S2. *Acropora millepora* genes with log2FC values and adjusted p-values for all timepoints compared to 0 h and predicted HSF1 binding sites in the 500bp region upstream of each gene.

Dataset S3. Reciprocal best blast hit results between species (14,431 genes identified) with log2FC values and HSF1 binding sites per species.

Dataset S4. Candidate biomarkers of thermal tolerance selected based on upregulation in *A. millepora* (log2FC ≥ 2, padj ≤ 0.001) but not in *G. fascicularis* (log2FC ≤ 1) at 3h of heat exposure, mimicking the HSF1 response.

