## Supplemental Information for "CRISPR/Cas9-mutagenesis reveals that varying dependence on HSF1 is associated with differences in coral heat tolerance"

**Materials and Methods**

***G. fascicularis* collection and husbandry**

In September 2021, four 640 L aquaria (“spawning systems”) (Coral Spawning International) were assembled as described by ref. 1. Each aquarium was filled with artificial seawater (ASW) (Red Sea Coral Pro Salt) and pre-conditioned with Marcorock, *Trochus* snails, and *Chaetomorpha* macroalgae. In October 2021, *G. fascicularis* adult colonies 8-15 cm in diameter were collected from the Great Barrier Reef, Australia, and shipped to Carnegie Science (Table S1). Each colony was mounted on a PVC platform using cyanoacrylate glue. Ten colonies were placed in System 1 and nine colonies were placed in System 2. Additional colonies were collected in February 2022 (distributed across all systems), September 2022 (placed into System 4), and November 2023 (placed into System 3) (Table S1).

All systems were maintained as described by ref. 1. Briefly, corals were fed Reef Brite Reef Enhance and freshly hatched *Artemia nauplii* five times per week with the circulation pump off for 35 min. Salinity was kept at 35 – 38 ppt using an automatic top-off pump to deliver reverse osmosis water. Calcium, magnesium, potassium, and other elements were added to each system at programmed intervals using a Dosatronic. Additional trace supplements were added after monthly water chemistry tests (Triton). Seawater dKH and pH were continuously monitored using an Alkatronic and nitrate, nitrite, and ammonia levels were checked weekly. Phosphate levels were stabilized using AL99 media (Unique Corals). Large debris was removed weekly by siphoning and 5-10% ASW changes occurred every 6-8 weeks.

**Shift of environmental parameters for laboratory-induced coral spawning**

In October 2021, all systems were set to follow an eight-year average temperature cycle (of non-sequential years; 1997-2017) of Moore Reef, central GBR, Australia, with a reduced maximum summer temperature of 28.5 °C (Figure 1B) as described by ref. 2. Solar and lunar photoperiod, sunrise time, sunset time, lunar intensity cycles, moonrise time, and moonset time were programmed with the parameters for Moore Reef in 2021 ^2^ using an Apex Pro controller as described by ref. 1. In November 2021, the environmental parameters were shifted independently in each aquarium to induce four spawning events per calendar year (Figure 1C). In System 1, one month of summer parameters was removed to shorten the gametogenic cycle to eleven months for predicted spawning in October 2022. After October 2022, System 1 was reset to induce a 12-month gametogenic cycle, with spawning predicted every October. In System 2, three months of summer parameters were removed to shorten the gametogenic cycle to nine months for predicted spawning in August 2022. After August 2022, System 2 was reset to induce a 12-month gametogenic cycle, with spawning predicted every August. In February 2022, System 3 was adjusted to add three months of spring parameters to extend the gametogenic cycle to fifteen months for predicted spawning in February 2023. After February 2023, System 3 was reset to induce a 12-month gametogenic cycle, with spawning predicted every February. Parameters in System 4 proceeded unchanged from the 12-month gametogenic cycle for predicted spawning in November 2022. After November 2022, six months of summer were removed to shorten the gametogenic cycle to six months, which predicted the next spawn in May 2023. After May 2023, System 4 was reset to induce a 12-month gametogenic cycle, with spawning predicted every May.

**Coral spawning, gamete collection, and controlled fertilizations**

Approximately four weeks prior to the predicted spawning event, the system’s internal clock was pushed forward eight hours so that sunset (all lights off) occurred in the morning. Spawning was checked between two and nine days post full moon from sunset until moonrise during both the month prior to predicted spawning and the month of predicted spawning. To control parental fertilizations, individual coral colonies were moved into 5.8 L isolation buckets before spawning began on predicted spawning days. Intact gamete bundles were collected within one minute of release using 5 mL transfer pipettes into 50 mL conical tubes. Spawning corals remained in isolation buckets until approximately one hour after their last gamete bundle was released and were then returned to the main tank.

To prepare gametes for fertilization, conical tubes were swirled to break up bundles. For female colonies, eggs were rinsed with a 125 µM filter sitting inside a 1 L food-safe bowl (Target, Room Essentials Plastic Cereal Bowl Polypro) filled with ASW. Eggs were gently rinsed with additional ASW. Most of the ASW was removed and eggs were placed into another 1 L bowl containing approximately 750 mL ASW. Eggs from each colony were kept in separate bowls in a 27 °C incubator with 12h: 12 h light: dark cycles at 25 µmol photons m^−2^ s^−1^ (Percival no. AL22L2C8). Every half hour, eggs were stirred lightly using a pipette to prevent adherence to the edge of the bowl. Gametes from male colonies were poured through a 125 µM filter to remove pseudo-eggs ^3^. Filtered sperm from individual coral colonies was transferred to a new conical tube and kept in a 27 °C incubator. Filtered sperm from all the male colonies was combined and diluted to a concentration of approximately 10^6^ cells mL^-1^. This bulk sperm mix was used for all subsequent fertilizations on that day.

Fertilizations were prepared by adding ~200 mL of ASW to a food-safe container (WebstaurantStore no. 128HD16COMBO). A pool of eggs from multiple females and ~6 mL of bulk sperm mix was added to the container, gently swirled, and kept at 27 °C. To estimate the success of fertilization after gametogenic cycle shifts (Figure 1D), multiple fertilizations (2-35 replicates per spawning event) were performed as described above with a known number of eggs (20-350 eggs per fertilization). Surviving larvae were counted at 24 hpf. For microinjections, fertilizations occurred at 1 h staggered intervals and were prepared as described above with ~200 eggs. Newly fertilized embryos were kept at 27 °C. Fertilizations continued for eight hours after initial spawning.

To assess optimal temperature for fertilizations and early development (Figure S1A), three replicate fertilizations were prepared in ASW at 23, 25, 27, 29, and 31 °C using temperature-controlled heating mats (Vivosun). For 20 °C, the three replicates were kept in a temperature-controlled room. ASW temperature was monitored continuously using a thermometer placed in a fourth container. Each replicate contained at least 60 eggs from multiple females and a standardized mixture of sperm from multiple males. Fertilizations were kept at each temperature until quantification after 24 h.

To assess optimal salinity for fertilizations and early development (Figure S1B), three replicate fertilizations were prepared in ASW at 31, 33, 35, 37, 41, and 43 ppt measured by a digital refractometer (Milwaukee Instruments) and placed in a 27 °C incubator. Each fertilization replicate contained at least 50 eggs from multiple females and a standardized mixture of sperm from multiple males. Fertilizations were kept at each salinity until quantification after 24 h.

To quantify the mean number of eggs per bundle over a shortened gametogenic length, bundles from a single female colony were collected during spawning in November 2022, May 2023, and May 2024 (Figure S1C). Bundles were collected unbroken from 6 – 12 polyps using a transfer pipette into individual 1.5 mL tubes. Each bundle was gently broken by pipetting and eggs were quantified using a stereo microscope.

An image time course of *G. fascicularis* embryo development was performed by fixing samples at known time intervals (Figure S1D). Samples in 100 µL ASW were lightly fixed by adding 100 µL of 4% PFA in ASW. All liquid was removed using a 25-gauge needle (BD PrecisionGlide) attached to a 1 mL syringe. Then, 500 µL of 4 % PFA in ASW was added and samples were kept overnight at 4 °C. The next day, the samples were washed 3 times in 500 µL of 1 x PBS and stored in 500 µL of 1 x PBS at 4 °C until imaging. Images were taken on a Leica MZ165 FC with a Flexacam C3.

To assess if the timing of first cell cleavage of zygotes varied as gametes aged, fertilizations were prepared as described above (Figure S1E). Staggered fertilizations between 0 and 9 h after gamete release contained eggs from multiple females and a standardized mixture of sperm from multiple males and were kept at 27 °C. Zygotes were checked using a stereo microscope every 15 - 20 min until the first cleavage furrow was observed.

To quantify if gamete age at time of fertilization impacted larvae survival over time (Figure S1F), staggered fertilizations occurred as described above. Survival measurements at 1, 9, 10, 11, and 15 hours after gamete release contain a single fertilization and the remaining time points each contain two fertilizations. Each fertilization contained at least 60 eggs from multiple females and a standardized mixture of sperm from multiple males. Larvae were scored for survival and given a full ASW change every day for four days.

**Synthesis, microinjection, and sorting of mTurquoise2 messenger RNA**

To generate capped messenger RNA of the Turquoise2 fluorescent protein (mTQ2) ^4^, a 50 µL PCR using KOD Hot Start polymerase (Sigma-Aldrich) was performed using 10 ng plasmid containing an SP6 promoter, mTurquoise2 sequence, and an SV40 poly(A) signal ^5^ (Table S8). The resulting 1242 base pair PCR product was confirmed for quality and size using a 1% agarose gel before digesting with DpnI [New England Biolabs (NEB)] for 60 min at 37 °C to remove the plasmid template. After incubation, the product was column purified using the DNA Clean & Concentrator - 5 (Zymo) and 100 ng of the resulting product was used in an overnight in-vitro transcription reaction using the HiScribe® SP6 RNA Synthesis Kit, following the manufacturer’s instructions (NEB). The next day, the transcription product was incubated with DNase-I for 35 min to remove the PCR template and RNA was precipitated with LiCl and washed twice with 70% ethanol following the manufacturer’s instructions. RNA quality and size was confirmed by running a 0.5 µL sample on a 2% agarose gel. A 5’ 7-methylguanylate cap (Cap-0) was added to 45 µg of RNA product using the Vaccinia Capping System (NEB) following the manufacturer’s instructions. The final mRNA product (Figure S2B) was column purified with an RNA Clean & Concentrator - 25 kit (Zymo). To obtain highly concentrated mRNA, 2 - 4 capping reactions were pooled and purified through a single column. All mRNA was stored at - 80 °C until injection.

An 8 µL mTQ2 mRNA injection mix was prepared in a 1.5 mL RNase-free tube with 3.52 µL (800 ng µL^-1^) mTQ2 mRNA, 1.0 µL 0.22-µM filtered 0.5% phenol red (MP Biomedicals), and 3.48 µL nuclease-free H_2_O. To control for injection, an 8 µL phenol-red-only injection mix was prepared in a 1.5 mL RNase-free tube as follows: 1.0 µL 0.22-µM filtered 0.5% phenol red and 7.0 µL nuclease-free H_2_O. Mixtures were freshly prepared on the injection day and kept on ice until injection. Microinjection proceeded as described in refs. 6 and 7, using the phenol red for visualization during injection.

At approximately 24 hpf, surviving larvae were counted (Figure S2D) and manually transferred to new ASW using a glass Pasteur pipette. For scoring, larvae were individually transferred to a 20 µL droplet of ASW on a glass slide and visually inspected under a Nikon E800 Eclipse compound microscope using a CFP filter (excitation bandpass 420 – 450 nm) with laser intensity at 60% (X-Cite, Excelitas Technologies) (Figure S2E). Larvae were visually scored as either having no expression (absence of cyan fluorescence) (Figure S2C; top), mosaic expression (Fig S2C; center), or uniform expression (Figure S2C; bottom). For imaging, a subset of larvae in each expression category were fixed in 4% paraformaldehyde and immediately imaged with the following parameters: 10x objective lens, scale: 8 – 13417, 60% laser, 100 ms exposure (MetaMorph, Molecular Devices) (Figure S2C). Prior to fixation, six larvae injected at the 1-cell stage with phenol red only (Figure S2C; top) and five larvae injected at the 1-cell stage with mTQ2 mRNA and scored as uniformly fluorescent (Figure S2C, bottom) were separated into new containers of ASW and kept at 27 °C in the dark to prevent spontaneous settlement. These larvae were live-imaged intermittently until 23 dpf to track cyan fluorescence over time (Figures S2F and S2G). Fluorescence in arbitrary units (a. u.) was determined using FIJI version 2.1.0/1.53c ^8^ with the ‘Measure’ tool to calculate the mean pixel value of each larva. Images shown in Figure S2C and Figure S2F were pseudo-colored with FIJI (version 2.14.0/1.54f) ^8^.

**DNA extraction and sequencing to generate a *G. fascicularis* genome**

To generate a megabase-scale genome assembly for *G. fascicularis*, coral bundles from an individual male coral colony were collected into a 50 mL centrifuge tube. The bundles were broken by gentle swirling and pseudo-eggs were removed using a 125 µL filter. Filtered sperm was divided into multiple 1.5 mL aliquots and were pelleted by centrifugation at 12,000 rcf for 5 min. The supernatant was removed and the dry pellet was flash frozen by placing the tubes in a mixture of dry ice and 100% ethanol and stored at -80 °C until processing. Two separate genomic DNA (gDNA) extractions were completed with two of the aliquots: one for high molecular weight (HMW) long-read Nanopore sequencing and the other for high-accuracy short-read Illumina MiSeq sequencing.

HMW gDNA for Nanopore sequencing was first extracted from ~15 µL of pelleted sperm. The Monarch® HMW DNA Extraction Kit for Tissue (NEB) and the Ultra-Long DNA Sequencing Kit (ULK; Oxford Nanopore Technologies) were used to extract HMW gDNA according to the manufacturer’s instructions, incorporating modifications introduced by the ULK protocol where relevant. The Pippin Pulse Power Supply (Sage Science), Sage Pulse Midi Gel Box (Sage Science), and Pippin Pulse software (v1.34) ^9^ were used for Pulsed-Field Gel Electrophoresis to visualize HMW gDNA samples before and after library preparation steps. The final library was loaded onto a Spot-ON R9 version flow cell (Nanopore) fitted into a Nanopore MinION sequencing device. MinKNOW software (v22.03.6) ^10^ was used for MinION sequencing using default parameters for the ULK protocol, with basecalling turned off. The runtime was set for 72 h. Sequencing was paused ~42 h into the run to wash the flow cell and load more library, according to the ULK protocol for optimizing yield with ultra-long viscous gDNA. To increase the number of long reads generated, a second run was completed as described above with a new flow cell.

Genomic DNA for high-accuracy Illumina MiSeq sequencing was extracted from a second aliquot of pelleted sperm following the protocol as described below, without the addition of glycogen. Sample quality was verified with an Agilent 2100 Bioanalyzer and DNA quantity was measured using the Qubit dsDNA BR Assay Kit (Invitrogen). The final genomic DNA library was prepared using the Illumina DNA Prep, Tagmentation kit and the Nextera DNA CD indexes (Illumina) following the manufacturer’s instructions with an input of ~400 ng gDNA. The final DNA library was run on an Illumina MiSeq with 300-bp paired-end reads.

**Evaluation and annotation of the *G. fascicularis* genome**

**Nanopore Base-calling and evaluating read lengths and qualities**

Basecalling of both nanopore runs was performed with GPU-enabled Guppy v6.1.2+e0556ff93 from the Microsoft Windows command prompt using super accurate mode (dna_r9.4.1_450bps_sup.cfg). Basecalled reads were analyzed with custom scripts to assess and visualize distributions of read lengths, predicted accuracy based on base-calling quality scores, and their relationships. The Guppy basecaller automatically partitioned the base-called reads into “pass” and “fail” folders using a mean Q-score cutoff of 10 that corresponds to an expected accuracy of 90%. As part of exploring the performance of different assemblers (detailed below), two different sets of the reads were used for each across a range of parameters: (i) all reads in both “pass” and “fail” folders, and (ii) just the reads in the “pass” folder. While the methods below focus on the results from using all reads or only the passing reads from both libraries combined, a similar number of assemblies was also initially generated from all reads or only the passing reads from each library separately. However, these were always outperformed (according to various metrics described below) by using both libraries (with otherwise the same assembler and parameters) and so are not detailed further.

**Assembling the Base-called Nanopore Reads**

The combined reads from both Nanopore libraries were used to generate hundreds of assemblies with 4 assemblers: 45 assemblies with Canu (version 2.2) ^11^, 7 assemblies with Flye (version 2.9-b1778) ^12,13^ 16 assemblies with Shasta (version 0.10.0) ^14^, and 504 assemblies with WtdBg2 (version 2.5) ^15^. For each read set (using all reads or just passing reads), we tried the default settings for each assembler as well as exploratory sweeps over various combinations of parameters, which are further detailed below. For all assemblers, where relevant, we set the expected genome size to 400 Mb. All assemblies were generated between June-September 2022. Consensus polishing modules from each assembler were used as default or as prescribed to produce the initial assemblies, although these aspects of the assembly pipelines were not considered as part of the post-assembly polishing rounds below.

For Canu (version 2.2), 45 assemblies were generated and evaluated. For all parameter sets described below, both default sensitivity and increased sensitivity parameterizations (corMhapSensitivity=high corMinCoverage=0 corOutCoverage=100) were tried. All assemblies had the following core set of parameters: gridOptions=--time=168:00:00 genomeSize=400m obtovlMemory=150g minMemory=16g corMemory=48g cnsMemory=20g minInputCoverage=5 stopOnLowCoverage=2 canuIterationMax=1 utgovlMemory=150g. Note that sometimes runs needed to be restarted from where it left off with even more resources, and the above parameters reflect the higher end of the resource needs. There was little guidance at the time (2022) on how to use Canu effectively with high-accuracy ultra-long nanopore reads with accuracies exceeding 95% and length N50 exceeding 100 kb (as we had). Therefore, four major approaches of how to treat the reads were explored: **(i)** Approach 1: treat as if they were standard raw nanopore reads (-nanopore -raw), which were previously considered error-prone; **(ii)** Approach 2: treat as if they were corrected nanopore reads (-nanopore -corrected); **(iii)** Approach 3: treat as if they were PacBio reads, which were considered less error-prone than nanopore at the time (-pacbio -raw); and **(iv)** Approach 4: treat with a specialized set of parameters for high accuracy, ultra-long Nanopore reads that used PacBio HiFi settings with various tweaks (described below in detail). For all four approaches, various parameter combinations described below were tried with each of the four following pairs of “minReadLength” and “minOverlapLength” parameters, respectively: (a) defaults, (b) 10 kb and 5 kb; (c) 20 kb and 10 kb; and (d) 30 kb and 15 kb. When treating reads as raw nanopore reads (approach 1 above), four pairs of error rate parameters were tried for “rawErrorRate” and “correctedErrorRate”, respectively: (a) defaults, (b) 0.30 and 0.075, (c) 0.20 and 0.050, and (d) 0.12 and 0.035. Similarly, when treating the reads as corrected nanopore reads (approach 2 above), four error rate parameters were tried for “correctedErrorRate” (note that “rawErrorRate” is irrelevant when using the -corrected flag): (a) defaults, (b) 0.30, (c) 0.20, and (d) 0.12. When treating the reads as if they were PacBio reads (-pacbio -raw) (approach 3 above), default error rates were used as specified by those flags. When treating reads as if they were ultra-long HiFi reads (approach 4 above), the following flags were used with all combinations of parameters: -pacbio-hifi 'batOptions=-eg 0.10 -sb 0.01 -dg 2 -db 1 -dr 3'. In addition to sweeping over different read sets (all or only passing), and different read and overlap lengths (described above), we either treated the reads as if they were still untrimmed (-untrimmed) or not (skip trimming) and tested two values of “correctedErrorRate” for all parameter combinations: 0.12 and 0.15. Note that these values are for reads with expected error rates of 6% and 7.5%, respectively, where overlap error rates for those reads are therefore expected to be double that.

For Flye (version 2.9-b1778), for each read set (all or only passing), the input reads were either classified as “--nano-hq” or “--nano-raw”, each with or without using the scaffolding parameter, “--scaffold”. Two other modes, “--meta” and “--keep-haplotypes”, we also tried in addition to the default assembly mode. However, as the special modes had little effect on the resulting assemblies, we ignored them further. All Flye assemblies were run with “--genome-size 400m”, and by default had one round of polishing (that is not considered part of the post-assembly polishing rounds below). Ultimately, we kept just 7 Flye assemblies to further evaluate.

Shasta (version 0.10.0) was used with four different configurations for each set of reads: Nanopore-May2022, Nanopore-Phased-May2022, Nanopore-UL-May2022, and Nanopore-UL-Phased-May2022. Although using 4 configurations on 2 read sets translates to 8 different assemblies, the phasing configurations generate three versions of the assembly: Haploid, Phased, and Detailed. All three versions of the assembly were evaluated in these cases, totaling 16 assemblies across both read sets. Note that older configurations (Nanopore-Oct2021, Nanopore-UL-Oct2021, Nanopore-UL-Jan2022) were also tested in an earlier phase of this project but not evaluated further after using the updated configurations released by the authors.

We generated 504 assemblies with WtdBg2 (version 2.5) as it was fast enough and light-weight enough to use nested for-loops over different values for a few parameters: subsampling k-mers with -S set to 4, 2, or 1; varying the minimum read depth for a valid edge with -e set to 3, 2, or 1; keeping or discarding “contained reads” by using -A or not; varying the minimum alignment length with -l set to 2048 (default), 10000, or 20000; and varying other options by trying available presets using -x in ont, preset1, preset2, preset3, or preset4 (noting that the “ont” preset allegedly maps to preset2 or preset3 depending on genome size). The wtpoa-cns consensus module was also subsequently run on all wtdbg2 assemblies.

Note that we also tested another assembler, Miniasm (version 0.3-r179) ^16^, in an earlier phase of this project using only the first Nanopore library. We generated 76 assemblies with Miniasm and Minimap2 (version 2.24-r1122) ^17^. For Miniasm, in addition to default settings, we tried combinations of skipping the second (-2) or both the first and second (-1 -2) read filtering steps, different coverage cutoffs (-c 1; -c 2; -c 5; -c 8), different required minimum overlap lengths (-s 10000 ; compared to 1000 default), and higher match minimums (-m 1000 ; compared to 100 by default) that seemed warranted in the ultra-long read setting. We also tried more stringent overlap accuracies (-i 0.5 ; -i 0.75 ; compared to default 0.05), prefiltering clearly contained reads (-R), less stringent small contig filtering (-e 1 ; -e 2) that might be warranted in a low-coverage ultra-long read setting, and increasing the maximal probing for bubble popping from the default of 50 kb to 5 Mb (-d 5000000). Nonetheless, Miniasm did not perform as well as the other assemblers (wtdbg2, Flye, Canu, Shasta), so was not further tested with both read sets.

**Selecting a candidate assembly from each assembler**

For genome size estimation and comparison, files associated with a previous reference genome assembly for *Galaxea fascicularis* (gfas_final_1.0) were downloaded from <http://gfas.reefgenomics.org/download/>: gfas_final_1.0.fasta.gz, gfas_1.0.transcripts.fasta.gz, gfas_1.0.proteins.fasta.gz, and gfas_1.0.genes.gff3.gz ^18,19^. A candidate assembly from each assembler was selected based on which had optimal contiguity statistics. Genome assembly contiguity across all assemblies generated was assessed with custom scripts (<https://github.com/JohnUrban/sciaratools2>) ^20^. For statistics that rely on an expected genome size, an expected genome size equivalent to the sum of all scaffold lengths in the previous assembly from reefgenomics.org, gfas_final_1.0, was assumed (334,165,880 bp). Assemblies with assembly sizes smaller than half the expected genome length were ignored. We focused on four contiguity metrics: the longest contig length, the NG50 contig length (the shortest contig length such that 50% of the expected genome size is assembled in contigs at least this long), LG50 (the number of contigs represented by the NG50 length), and expected contig lengths all normalized to the same denominator (expected genome size) rather than using individual assembly sizes ^21^. The highest ranking assembly for each assembler were selected based on ranking them by favoring higher maximum contig lengths, higher NG50s, lower LG50s, and higher normalized expected contig lengths. The parameters for the selected assemblies were as follows.

The selected Canu assembly was from using all Nanopore reads (not just the “pass” reads) and treating them as if they were PacBio HiFi reads (--pacbio-hifi) with tweaked bat options ('batOptions=-eg 0.10 -sb 0.01 -dg 2 -db 1 -dr 3'), using default sensitivity settings, default minimum read and overlap length settings, a correctedErrorRate of 0.12, and skipping the trimming step (not using the “-untrimmed” flag):

canu -p hifi-CE0.12 -d hifi-CE0.12 gridOptions="--time=168:00:00" genomeSize=400m obtovlMemory=48g minMemory=16g corMemory=48g cnsMemory=50g minInputCoverage=5 stopOnLowCoverage=2 canuIterationMax=1 utgovlMemory=32g 'batOptions=-eg 0.10 -sb 0.01 -dg 2 -db 1 -dr 3' -pacbio-hifi reads.fasta correctedErrorRate=0.12

For Flye, the selected assembly was produced using only the passing reads treated as high quality nanopore reads (--nano-hq) and with the scaffolding flag set (--scaffold):

flye --scaffold --genome-size 400m --nano-hq ${READS} --out-dir results --threads ${P}

For Shasta, the selected assembly was produced using all of the reads and the “Nanopore-May2022” configuration. As the reads used were “ultra long” with read N50s exceeding 100 kb, it was surprising that ultra-long parameters did not produce the most contiguous assembly.

For WtdBg2, the selected assembly was produced using all of the reads and by adopting “preset3”, requiring alignments to be at least 20 kb long (-l 20000), keeping contained reads during alignment (-A), using “-S 2” to control the amount of kmers subsampled for indexing, and a minimum read depth of 2 for a valid edge (-e 2).

wtdbg2 -x preset3 -g 400m -t 8 -i reads.fastq -fo asm -S 2 -e 2 -A -l 20000

wtpoa-cns -t 8 -i asm.ctg.lay.gz -fo asm.ctg.fa

**Initial Long-Read Polishing of the representative assemblies from each assembler**

Assemblies were polished using the Nanopore reads with seven different pipelines, using all reads (not just passing reads) in each pipeline. Earlier testing with just one Nanopore library showed similar or sometimes inferior results when using only passing reads. Thus, all reads were used moving forward. In total, 13 different polished assemblies were produced for each input assembly. All polishing was performed circa October 2022 using the then most recent releases of relevant software. All pipelines described below were run using racoonFlyFish (<https://github.com/JohnUrban/racoonFlyFish>), which conveniently automates multi-round genome assembly polishing with long reads using combinations of RaCon ^22^, Flye ^12^, and Medaka (<https://github.com/nanoporetech/medaka>). Overall, there were 13 long-read polished outputs across 7 pipelines kept for evaluations, as described below.

Pipeline 1 consisted of using Medaka (medaka_consensus version 1.7.2) from Oxford Nanopore Technologies, using the model most relevant to our base-called reads available at the time (r941_min_sup_g507).

Pipeline 2 consisted of using the polishing module from Flye (version 2.9-b1778) to do 4 rounds of polishing. The polished assembly from each round was kept for evaluations, resulting in 4 flye-polished assemblies for each input assembly. Non-default parameters for Flye polishing were “--genome-size 400m --nano-raw --iterations 4”.

Pipeline 3 consisted of using RaCon (version 1.5.0) with Minimap2 (version 2.24-r1122) to do 4 rounds of polishing. Again, the polished assembly from each round was retained for evaluating, resulting in 4 RaCon-polished assemblies for each input assembly.

Pipeline 4 consisted of running Medaka on the assembly produced after the fourth round of Flye polishing.

Pipeline 5 consisted of running Medaka on the assembly produced after the fourth round of RaCon polishing.

Pipeline 6 consisted of running one round of Flye polishing followed by one round of RaCon polishing.

Pipeline 7 consisted of running one round of Flye polishing, then one round of RaCon polishing, and finished with one round of Medaka polishing.

**Further evaluations of the candidate assemblies from each assembler and their 13 polished counterparts**

After long-read polishing, there were 14 representative assemblies for each assembler: the input and 13 polished counterparts. Allowing for the possibility that assemblies had higher quality consensus sequences before long-read polishing, all 14 representative assemblies corresponding to each assembler were considered as substrates for further evaluations to select a final representative for each assembler. The assemblies were evaluated using metrics focused on gene completeness, sequence completeness, and consensus accuracy. To approximate gene completeness, two programs were used: BUSCO ^23^ and LiftOff ^24^. For both BUSCO and LiftOff, assembly ranking favored fewer missing genes. To estimate sequence completeness and consensus accuracy, k-mer based and read mapping approaches were taken, both using the high accuracy Illumina MiSeq data. For both the k-mer and read mapping approaches, assembly ranking favored higher completeness scores and higher QV estimates, the latter of which are Quality Value estimates of the assemblies corresponding to -10*log10 (estimated error rate). All metrics and interpretations are further detailed below.

BUSCO searches for genes that are present as single-copy orthologs in the majority of species of a given clade and are therefore expected to be in the current assembly as single-copy genes. It reports the percent missing and percent found (also noting whether they were found complete or fragmented and whether they were single-copy or multi-copy). We used BUSCO’s clade detection feature to pick gene groups for evaluations (-m geno --auto-lineage-euk). BUSCO gave reports for Eukaryota (255 genes) and Metazoa (954 genes). We confirmed that BUSCO offered no gene groups for clades closer to the genus or species level (<https://busco.ezlab.org/list_of_lineages.html>).

LiftOff is a tool to lift gene annotations (or any feature described by a GFF) from one genome assembly over to another. In doing so, it can also report the number of annotations it was unable to map, which can be interpreted as “missing genes”. Files associated with the previous reference genome assembly for *Galaxea fascicularis* (gfas_final_1.0) were downloaded from <http://gfas.reefgenomics.org/download/>: gfas_final_1.0.fasta.gz, gfas_1.0.transcripts.fasta.gz, gfas_1.0.proteins.fasta.gz, gfas_1.0.genes.gff3.gz. Using LiftOff, we lifted over the GFF file featuring gene annotations generated for Gfas_final_v1.0 to each de novo assembly. Thus, we were able to evaluate the presence or absence of 22,418 previous gene annotations corresponding to ~171 Mb of gene sequence.

For the k-mer based approach to estimating sequence completeness and consensus accuracy, GenomeScope2 (version 2.0) ^25,26^ and Merfin (merfin snapshot v1.1-development +6 changes, r200 fc4f89a3d62504dece52f5cac22448c3669c7f1c) ^27^ were used, the latter of which descends from Merqury ^28^ and relies on a provided version of Meryl ^28^. Briefly, Meryl was used to count, get the union-sum of, and produce a corresponding histogram for kmers in the reads for FastQ files corresponding to both Illumina MiSeq mates. The kmer histogram was used with Genomescope2, using the fitted option, to get model parameters (the kcov peak and look-up table). Note that GenomeScope2 provided a heterozygosity estimate of 0.0204 (i.e. ~2.04%). Merfin “hist” and “completeness” was then used with the meryl kmer database and genomescope2 model parameters to evaluate each assembly. Ultimately, this provided three metrics: k-mer based completeness, the Merqury QV, and the Merfin QV. The Merqury QV only considers kmers in the assemblies that are missing from the reads whereas the Merfin QV considers both those missing kmers as well as kmers that are overrepresented in the assemblies compared to the reads.

For the read mapping approach, the Illumina MiSeq mates (R1 and R2) were mapped independently with Minimap2 (version 2.24-r1122) using the following parameters: -a -x sr --end-bonus 5. Completeness corresponds to the percent of reads that mapped. Consensus accuracy is estimated from the SAM formatted output using a python script that tracks the alignment information for each read returning the number of matches, mismatches, deletions, insertions, and unaligned bases. This information is used to compute two average error rates: (i) (1 - m/(m+mm+d+i)), and (ii) (1 - m/(m+mm+d+i+u)); where ‘m’, ‘mm’, ‘d’, ‘i’, and ‘u’ stand for the number of matches, mismatches, deletions, insertions, and unaligned bases, respectively, summed across all reads. The first error rate accounts for only alignment blocks whereas the second adjusts the estimate by accounting for both alignment blocks and unaligned bases. The error rates are the sum of three sources of “error”: heterozygosity rate, base-calling error, and assembly error. Although the raw error rates were sufficient to compare assemblies since the first two are constant in the read set, the error rates could be adjusted to estimate the assembly error rate by subtracting both the estimated heterozygosity rate learned with GenomeScope2 (0.0204) and the base-calling error rate learned from base quality scores, which leaves only the error from each given assembly. The average read error was estimated based on the distribution of all base-calling quality scores in the FastQ file, obtained using Seqtk (<https://github.com/lh3/seqtk>): seqtk fqchk -q 0. When estimating assembly error rates by subtracting out the effect from heterozygosity and read error, a QV of 60 was given to anything with an assembly error rate of 0.000001 or lower. Assemblies that maximize the number of read alignments and minimize the error estimates above were ranked higher than those with fewer alignments and/or higher error rates. Error rates were converted to QVs as above to compare with kmer-based QVs.

Ultimately, assemblies were ranked based on minimizing (i) the number of missing Eukaryotic BUSCOs, (ii) the number of missing Metazoan BUSCOs, and (iii) the number of unmapped Gfas1 genes, as well as maximizing (i) Merfin K-mer-based Completeness, (ii) R1 Map-based Completeness, (iii) R2 Map-based Completeness, (iv) Merqury QV, (v) Merfin QV, (vi) R1 Alignment QV, (vii) R1 Adjusted QV, (viii) R2 Alignment QV, (ix) R2 Adjusted QV. Taking average ranks various ways all gave the same results.

For all four assemblies, long read polishing did improve the completeness and accuracy metrics compared to their pre-polished counterparts. However, different pipelines worked best for each. For each assembler, the assembly (out of 14) that appeared to optimize the evaluation results was chosen for further polishing with the high accuracy short reads using Pilon (see next section). The long-read polishing that worked best is as follows:

- 4 rounds of Flye-polishing followed by Medaka for the selected Canu assembly;
- 2 rounds of Flye-polishing for both the selected Flye and WtdBg2 assemblies;
- 4 rounds of Flye-polishing for the Shasta assembly.

**High accuracy short read polishing of selected assemblies**

The four selected long-read polished assemblies were polished using the high accuracy Illumina MiSeq reads with Pilon (version 1.24) ^29^. In addition, their pre-long-read-polishing counterparts were also polished with Pilon to allow the possibility that skipping long-read polishing produced better completeness and sequence accuracy. Thus, there were two assemblies corresponding to each of the four assemblers, totaling 8 assemblies for short read polishing with Pilon.

In an earlier phase of this project, using only a single Nanopore library, we found that the consensus sequence accuracy was high (>99%) directly from the assembler prior to any polishing, and we found that being conservative with short-read polishing gave higher quality results. That is, as opposed to being aggressive with short read polishing by using all mapped reads (MAPQ>0) and bases of any quality (BQ>0), better results were achieved by using (i) using Bowtie2 ^30^ in --very-fast mode (as higher sensitivity modes seemed to more aggressively make erroneous changes) and specifying a maximum fragment size of 600 bp, (ii) using the end-to-end mapping mode (as opposed to allowing local alignments), (iii) using only concordantly mapped reads that were confidently mapped (MAPQ>10 or >30) for polishing (not orphaned, discordantly-mapped, or more ambiguously-mapped reads),  (iv) using only high quality base calls within the confidently mapped reads (BQ>10 or >30), and (v) ignoring the 10 bp at each end of a read alignment. These principles were retained for the final short-read polishing phase of this project as described below.

For each of the 8 assemblies, three rounds of Pilon polishing were run with two pairs of differential parameters for Pilon (--minmq and --minbq): (i) conservative filtering (MAPQ and BQ > 10), and (ii) very conservative filtering (MAPQ and BQ > 30). Otherwise, other parameters for Pilon were held constant (--changes --vcf --diploid --nostrays –fix bases –flank 10). For round 1, reads were mapped to the initial input. For each subsequent round, reads were re-mapped to the output polished assembly from the previous round. For each round, reads were mapped with bowtie2 (--maxins 600 --end-to-end -q --very-fast), filtered for MAPQ and concordant mapping with SAMtools ^31^ “view”, sorted with SAMtools “sort”, and duplicates marked and removed with Picard MarkDuplicates (<http://broadinstitute.github.io/picard/>) ^32^. Polished assemblies from each round were retained for further evaluation to allow the possibility that fewer rounds were better. In total, 6 different short-read polished assemblies were produced for each input assembly. All polishing was performed circa October 2022 using the then most recent releases of relevant software. All pipelines were run using AutoPilot (<https://github.com/JohnUrban/AutoPiloT>), which conveniently automates multi-round genome assembly polishing with Pilon. The assemblies from all short-read polishing stages were evaluated as above for long-read polished assemblies, using the same metrics focused on gene completeness, sequence completeness, and consensus accuracy. In all cases, Pilon polishing moderately improved upon long-read polishing, and starting with the long-read polished intermediates did better than skipping long-read polishing. For all four long-read polished assemblies, using 3 Pilon rounds with filtering for MAPQ and BQ > 10 performed best. Thus, the four candidates selected at this stage of the workflow were each polished by long reads followed by short reads.

**Purging duplicates, final polishing, and final evaluations of the assemblies**

The four long-read polished, short-read polished assemblies were selected for purging duplicates and haplotigs with purge_dups ^33^. This step separates the assembly into a Primary assembly and an Associated assembly. The Primary assembly contains a single representation of all loci through the genome. The Associated assembly contains alternative haplotypes on smaller contigs that map completely inside a contig from the primary assembly. For all four assemblies, the size of the Associated assembly rivaled the size of the Primary assembly. This was due to the high accuracy of the Nanopore reads and parameterization of the assemblers, which produced nearly diploid assemblies as opposed to a single haploid representation. This diploid representation meant that mapping short reads would have looked ambiguous for the majority of loci in the previous polishing step. Therefore, for each of the four assemblies, their Primary and Associated counterparts were subject separately to three more rounds of Pilon polishing. In addition, we also tried three rounds on both of them still together for comparison. Evaluations were run on the output of the third Pilon rounds.

After a final round of evaluations focusing on gene completeness, sequence completeness, and consensus accuracy, the assembly that was selected as the final assembly for gene annotation was the one produced by Canu when treating the reads somewhat like ultra-long PacBio HiFi reads, long-read polished with 4 rounds of Flye-polishing followed by Medaka, short-read polished with 3 rounds of Pilon using MAPQ and BQ > 10, purged with purge_dups, and is primary and associated counterparts subject independently to more 3 rounds of Pilon again using MAPQ and BQ > 10. This assembly was named Carnegie_Gfas_genome_v1.0.

**Gene Annotation**

For the purpose of soft-masking repeat sequences for the gene annotation pipeline, we developed a de novo, species-specific repeat sequence library, Carnegie_Gfas_repeat_library_v1, from the Primary contigs of the final selected assembly using RepeatModeler version 2.0.3 ([http://www.repeatmasker.org](http://www.repeatmasker.org/)) ^34^. The Primary and Associated genome assemblies were kept separate to produce Primary- and Associated- gene annotation sets independently. Each partition was subject to the annotation pipeline as described below.

Prior to gene annotation, repeats were identified and soft-masked using RepeatMasker version 4.1.2-p1 (<http://www.repeatmasker.org>) ^35^, using NCBI/RMBLAST version 2.11.0+ (<https://www.repeatmasker.org/rmblast/>) ^36,37^ with the species-specific repeat library described above (Carnegie_Gfas_repeat_library_v1). As briefly described next and detailed further below, gene annotation was performed similar to the TSEBRA protocol ^38^ described for combining Braker1 results trained with RNA-seq data ^39^ with Braker2 results trained with protein data ^40^. However, after selecting gene models with TSEBRA, we ran an additional iteration of Braker2 with an updated protein dataset that included the initial TSEBRA outputs as well as predicted proteins from reference-guided transcriptome assemblies produced from reads mapped with HiSat2 ^41^ followed by assembly with StringTie ^42^ or PsiClass ^43^, transcript sequence extraction with Gffread ^44^ and protein sequence prediction with Borf ^45^. TSEBRA was then used again with the initial Braker1 RNA-seq-based gene models and the second iteration of Braker2 gene-models from the updated protein evidence that included proteins based on the RNA-seq data. The different stages of this pipeline are described in more detail below.

Step 1: Gene models from RNA-seq trained Braker 1

For the input to Braker1, STAR ^46^ was used to independently map single-end 75 bp strand-specific RNA-seq reads from 216 different samples, equating to over 1 billion reads. Specifically, STAR was used to index either the Primary or the Associated genome assembly separately, and subsequently to map all reads to that index with the following parameters: --twopassMode Basic --readFilesCommand zcat --outSAMtype BAM SortedByCoordinate. The sorted BAMs from all 216 samples were then merged with SAMtools. SAMtools was then used to partition the merged BAM into two files: reads mapped to the forward-strand (-F 16) and reads mapped to the reverse-strand (-f 16) with the SAMtools view sub-command. Braker1 (Braker version 2.1.6) was run using the merged, strand-separated BAMs (--species gfas_braker1 --genome=${ASM} --UTR=on --softmasking --stranded=+,- --bam=${FWD},${REV} --workingdir=braker-UTR --cores=16). Although we set out to predict UTRs (--UTR=on), we found that the gene models with UTRs (augustus.hints_utr.gtf) were very poor compared to the gene models predicted without UTRs (augustus.hints.gtf) on the same run as a prior part of the pipeline leading up to UTR prediction. For example, the models without UTR training (augustus.hints.gtf) looked very similar to the gene models produced by Braker2 with only protein evidence and very similar to the gene models from StringTie and PsiClass reference-guided transcriptome assemblies, whereas the UTR-trained models (augustus.hints_utr.gtf) were often very different and in their own category. In fact, the Braker1 output messages from annotating the Primary assembly reported that “UTR training decreased AUGUSTUS ab initio accuracy” from 0.81 in the initial training without UTRs to 0.47 after UTR training. Therefore, we used only the no-UTR gene models (augustus.hints.gtf) going forward and discarded the UTR-trained models.

Step 2: Gene models from protein-trained Braker 2

To use Braker2 with protein data, the initial protein set for iteration 1 included >1.5 million metazoan proteins from OrthoDB v11 ^47^; 22,418 proteins from a previous *Galaxea fascicularis* gene annotation set produced on a highly fragmented genome assembly ^18,48^; and the 8 sets of proteins available for the taxonomic Class, Hexacorallia, on NCBI, which included 3 sea anemones and 5 stony corals, totaling 255,861 proteins, each species ranging from ~25-42 thousand: *Nematostella vectensis* (sea anemone, GCF_000209225.1, 34311 proteins), *Exaiptasia diaphana* (sea anemone, GCF_001417965.1, 27753 proteins), *Actinia tenebrosa* (sea anemone, GCF_009602425.1, 27037 proteins), *Acropora digitifera* (stony coral, GCF_000222465.1, 33878 proteins), *Orbicella faveolata* (stony coral, GCF_002042975.1, 32587 proteins), *Stylophora pistillata* (stony coral, GCF_002571385.1, 33252 proteins), *Pocillopora damicornis* (stony coral, GCF_003704095.1, 25183 proteins), and *Acropora millepora* (stony coral, GCF_013753865.1, 41860 proteins). The NCBI datasets were downloaded from <https://www.ncbi.nlm.nih.gov/data-hub/genome>, specifying “Hexacorallia” and filtering for annotated reference genomes: <https://www.ncbi.nlm.nih.gov/data-hub/genome/?taxon=6102&reference_only=true&annotated_only=true&refseq_annotation=true&genbank_annotation=true>.

Braker2 (Braker version 2.1.6) was run with this initial protein set (--species gfas_braker2 --genome=${ASM} --prot_seq=${PROTEINS} --softmasking --cores=16 --workingdir=braker-prot).

Step 3: TSEBRA selection of gene models from Braker1 and Braker2 (iteration 1).

TSEBRA ^38^, was used to select a final set of gene models from the Braker1 and Braker2 outputs given all of the evidence (-g ${BRAKER1}/augustus.hints.gtf,${BRAKER2}/augustus.hints.gtf -c default.cfg -e ${BRAKER11}/hintsfile.gff,${BRAKER2}/hintsfile.gff -o brakerTsebra_combined.gtf --score_tab tsebra-scores.tab). The TSEBRA helper script, rename_gtf.py, was used to rename the TSEBRA-selected gene models. Gffread ^44^ was used to extract transcript and protein sequences from the genome FASTA using the GTF file with renamed TSEBRA-selected gene models.

Step 4: Creating an updated protein database with the addition of RNA-seq-informed proteins.

Briefly, the protein dataset was expanded as follows:

(1) The above Braker1/Braker2/TSEBRA pipeline (Steps 1-3) was done separately on the Primary and Associated assemblies. The resulting TSEBRA-selected proteins from each assembly, which contain RNA-seq-based Braker1 models, were added to the initial protein set.
(2) In addition, as described in detail below, two reference-guided transcriptome assemblies were generated for each the Primary and Associated assemblies (4 transcriptomes total), and the predicted proteins from all 4 transcriptome assemblies were added to the initial protein set.

The following was done to obtain transcriptome-based protein sequences. HiSat2 ^41^ was used to index each partition of the assembly (Primary and Associated), and to subsequently map the reads from all 216 strand-specific RNA-seq samples separately to each (-x index -U R1.fastq.gz --rna-strandness R --dta -p 16). HiSat2-mapped reads were then assembled using two different pipelines: StringTie ^42^ and PsiClass ^43^. StringTie was first run separately on each sample BAM file to assemble transcripts of at least 200 nucleotides in length (--rf -m 200 -p 16 -l sampleID -o sampleID.gtf reads.bam), then subsequently used to merge all 216 individual assemblies (--merge -p 4 -o stringtie-merged.gtf -m 200 -l gfas-<assembly>-MSTRG mergelist.txt, where <assembly> was either ‘primary’ or ‘associated’). PsiClass also produced transcriptome assemblies from each sample as well as a single merged/voted assembly from all 216 samples, but with a single command (psiclass -p 16 --lb bam.fofn --stranded rf; where bam.fofn is a file-of-file-names with the absolute file path to all 216 BAMs). Gffread was used to extract the transcript sequences from the multi-sample transcriptome assembly from each assembler (not the assemblies from individual samples), yielding two transcriptomes (1 StringTie and 1 PsiClass) for each the Primary and Associated genome assemblies (4 transcriptomes total). Borf ^45^ was used to predict protein sequences of at least 70 amino acids from each set of strand-specific transcripts (borf -l 70 transcripts.fa). These borf-predicted transcriptome proteins were also added to the updated protein database.

Step 5: Gene models from Braker2 with the RNA-seq-informed updated protein database.

Braker2 was then run again with the updated protein set (created in Step 4) that ultimately contained:

(i) The entire initial protein dataset (from Step 2 above),
(ii) TSEBRA-selected proteins from the Primary assembly (from Step 3 above),
(iii) TSEBRA-selected proteins from the Associated assembly (from Step 3 above),
(iv) Borf-predicted proteins from the Primary StringTie transcriptome (from Step 4),
(v) Borf-predicted proteins from the Associated StringTie transcriptome (from Step 4),
(vi) Borf-predicted proteins from the Primary PsiClass transcriptome (from Step 4), and
(vii) Borf-predicted proteins from the Associated PsiClass transcriptome (from Step 4).

Thus, in addition to the initial protein dataset, the RNA-seq data was able to influence the training of new Braker2 gene models via protein evidence from:

(1) the protein sequences from TSEBRA-selected Braker1 gene models produced by training on STAR-aligned strand-specific RNA-seq reads, and

(2) additional protein sequences predicted by Borf from transcriptome assemblies created from HiSat2-aligned strand-specific RNA-seq reads, StringTie, and PsiClass.

Where there were disagreements between Braker1 gene models trained using RNA-seq (Step 1) and the first iteration of Braker2 gene models trained with the initial protein set (Step 2), we found that the Braker1 models had higher concordance than Braker2 with transcript models produced from StringTie and PsiClass. This result is consistent with Braker1 being trained with the same RNA-seq data used to create the transcriptomes, and with the first iteration of Braker2 (Step 2) being “blind” to the RNA-seq data. However, in the set of the gene models from the second iteration of Braker2 (Step 5) that used the updated protein database, there were far fewer disagreements with Braker1 and transcriptome gene models, which would be expected as the updated training set contained proteins from Braker1 and the transcriptomes.

Step 6: TSEBRA selection of gene models from Braker1 and Braker2 (iteration 2).

TSEBRA was used one last time to combine gene models from Braker1 trained with RNA-seq data (Step 1) and from the second iteration of Braker2 trained with the updated protein set (Step 5). This produced a final gene set for each the Primary and the Associated assemblies. As above, rename_gtf.py was used to rename the TSEBRA-selected gene models. Gene identifiers from the Primary or Associated annotations have the following prefix: carnegie_gfas-primary-1.0 or carnegie_gfas-associated-1.0. Again, Gffread was used to extract transcript and protein sequences from the corresponding GTF and genome assembly files. This TSEBRA-selected set of gene models constituted our final official gene set (OGS), which was named carnegie_gfas_primary_OGS_v1.0 for the Primary assembly (and carnegie_gfas_associated_OGS_v1.0 for the Associated, which was processed separately).

Braker3 note:

An early version of Braker3, which was optimized to train gene models on RNA-seq and protein evidence at the same time as opposed to separately in the TSEBRA pipeline, was available around the time of finishing this work (early 2023), which also makes use of StringTie transcriptome assemblies to inform gene model training. To be thorough, we also ran Braker3 and compared the Braker3 gene models to those produced by our modified Braker1/Braker2/TSEBRA pipeline above. Our pipeline had fewer missing Metazoan BUSCOs than the Braker3 results, perhaps because our approach leans towards sensitivity (finding more real genes, but potentially also more false positives) while Braker3 may lean towards specificity (finding fewer false positives at the expense of more false negatives). For example, our modified Braker1/Braker2/TSEBRA pipeline yielded more genes (42,035 vs 26,101) and more proteins than Braker3 (48,022 vs 29,781) on the Primary assembly. Otherwise, we found that the majority of shared gene models between them were either the same or similar. Where there were disagreements, the Braker3 models seemed to merge neighboring genes that were kept as separate genes in our pipeline. Braker3 uses only StringTie to produce a merged transcriptome assembly to predict proteins from. In our work, we noticed that the StringTie assemblies had a higher propensity of creating long gene models, seemingly from merged neighboring genes, than gene models from PsiClass, Braker1, and Braker2. Thus, the reliance on StringTie alone in Braker3 may have biased it toward the longer, possibly-merged gene models. Given these observations, we chose to continue using the gene models produced from our modified Braker1/Braker2/TSEBRA pipeline.

**Functional Annotation**

Proteins from our modified Braker1/Braker2/TSEBRA gene annotation set were functionally annotated using BLASTP ^36^, InterProScan ^49^ and Google’s ProteInfer ^50^. BLASTP was used to find all hits (-word_size 3 -evalue 1e-2) for all proteins in the following BLAST databases in March 2023: (i) Non-redundant UniProtKB/SwissProt ^51^ protein sequences (ftp://ftp.ncbi.nlm.nih.gov/blast/db/swissprot.tar.gz); (ii) NCBI Landmark database (ftp://ftp.ncbi.nlm.nih.gov/blast/db/landmark.tar.gz); (iii) OrthoDB v10 ^52^; (iv) NCBI RefSeq ^53^ protein database (ftp://ftp.ncbi.nlm.nih.gov/blast/db/refseq_protein.*.tar.gz); and (v) NCBI Non-Redundant (nr) protein database (ftp://ftp.ncbi.nlm.nih.gov/blast/db/nr.*.tar.gz). The best BLASTP hit for each protein for the results from a given database was identified as the hit with the highest bitscore, and a more stringent set of best hits was created by filtering out hits with e-values higher than 0.0005 or bitscores lower than 50. For the InterProScan analysis, all protein sequences were further characterized using the InterPro ^54^ protein family and domain database and InterProScan version 5.56-89.0 (-dp -iprlookup -goterms -f tsv,xml,json,gff3, ^55^. All available analyses were executed: CDD, Coils, Gene3D, Hamap, MobiDBLite, PANTHER, Pfam, Phobius, PIRSF, PIRSR, PRINTS, ProSitePatterns, ProSiteProfiles, SFLD, SignalP, SMART, SUPERFAMILY, TIGRFAM, TMHMM. When necessary, software licenses were obtained (SignalP, Phobius, TMHMM). Finally, Google’s ProteInfer was run on all proteins in March 2024 with the then latest-available version using a low reporting threshold of 0.01, which allows filtering the output for higher confidence thresholds such as the default of >0.5, but also allows one to see lower confidence results if desired: python3 proteinfer.py -i ${FASTA} -o ${TSV} --reporting_threshold 0.01.

**Design of heat-stress experiments for RNA sequencing**

*A. millepora* colonies were collected from the Great Barrier Reef (GBR; Palm Island Reef) and kept in flow-through aquaria at the National Sea Simulator Facility at the Australian Institute of Marine Science, Queensland, Australia. Coral colonies spawned in November 2023 and after washing the gametes (as described in ref. ^6^), bulk fertilizations from multiple parental colonies were performed. *G. fascicularis* gametes were fertilized as described above following a 12-month gametogenic cycle. Developing larvae for each species were kept in polypropylene containers at 27 °C with 12h: 12 h light: dark cycles at 25 µmol photons m^−2^ s^−1^s and water-changed daily until experimentation.

The following heat-stress experiment was conducted for each species independently. At 3 dpf, three replicates of 40 larvae were added to individual wells in 6-well plates containing 7 mL of seawater and were placed into a 34 °C incubator. Seawater temperature was continuously monitored using a submersible data logger (HOBO pendant; Onset Computer Corp). At 0, 1.5, 3, 6, 12, and 24 h, larvae replicates were collected and placed in individual 1.5 mL tubes. All ASW was removed with a 0.5 x 40 mm needle (BD) and syringe, and 1 mL of nucleic acid preservation buffer ^56^ was added. Samples were placed at 4 °C overnight and then moved to -20 °C until processing.

Total RNA was extracted as described in ref. ^57^, with some modifications. Briefly, the 40 larvae were placed into a new sterile 1.5 mL centrifuge tube containing 100 µL of TRIzol reagent (Ambion). The larvae were homogenized with an RNase-free plastic pestle for 20 sec and then an additional 900 µL TRIzol and 100 µL of 0.5 mm glass beads (BioSpec Products) were added to each tube. The samples were vortexed on high for 20 sec with the tube on its side and then 10 µL of RNA-grade glycogen (Thermo Scientific) was added to each tube. Samples were incubated for 10 min at RT. Next, 100 µL of bromochloropropane (Sigma-Aldrich) was added to each tube which were then vortexed vigorously for 30 sec and incubated again for 10 min at room temperature. Samples were centrifuged at 16,000 rcf for 15 min at 4 °C. Five hundred microliters of the upper aqueous layer were transferred to a new 1.5 mL tube containing 250 µL High-Salt Solution for Precipitation (Plant) (Takara Bio) and 250 µL 100% isopropanol and stored at -20 °C overnight. Following precipitation, the samples were pelleted by centrifuged at 16,000 rcf for 15 min at 4 °C. The supernatant was discarded, and the pellet was washed with 1 mL of 75% ice-cold ethanol twice with a centrifugation step of 7,600 rcf for 5 min at 4 °C in between washes. After removing the supernatant and allowing the pellet to air dry for 1 min, the sample was resuspended in 50 µL nuclease-free H_2_O. Each sample was DNase-I treated and purified through a Zymo Quick RNA Microprep Plus Kit following the manufacturer's instructions and eluted in 15 µL nuclease-free H_2_O at 65 °C. The 15 µL eluate was passed through the purification column a second time. RNA samples were stored at -80 °C until library generation.

**RNA sequencing and analysis**

The quality and quantity of RNA for each sample was determined using an Agilent 2100 Bioanalyzer. RNA libraries were prepared using the Illumina TruSeq Stranded mRNA Kit (Illumina) and TruSeq RNA CD Index Plate (Illumina) following the manufacturer’s instructions with an input of ~400 ng of RNA from each sample. Pooled libraries of indexed samples were run on an Illumina NextSeq 500 using 75-bp single-end reads and 8 x 8 bp dual indexing. Two independent sequencing runs (all samples were included in each run) yielded between 6.8 and 18 million reads per sample, with a median of 13.06 and 12.49 million reads per sample for *A. millepora* and *G. fascicularis* samples, respectively.

RNA-seq data was aligned to either the published *A. millepora* genome ^58^ or the newly assembled *G. fascicularis* genome (this study) following the nf-core/rnaseq (version 3.12.0) bioinformatic pipeline ^59,60^. Salmon pseudo-count files were used as input for DESeq2, using the *tximport* function to account for transcript length for all subsequent analyses. DESeq2 was run with default parameters ^61^ for each coral species independently. Normalized count matrices obtained with DESeq2 median of ratios were used for all heatmaps, these values were then normalized based on the mean expression at time 0 h for each species separately. Genes involved in the UPR (Figure 2G) were manually curated by selecting genes that have been described as part of the unfolded protein response ^62–65^ (KEGG Pathway: hsa04141) and including orthologous genes that were differentially expressed in at least one of the species at any time point. For the frontloading analysis and *HSF1* normalized expression comparisons, TPM count matrices obtained from nf-core/rnaseq version 3.12.0 analysis pipeline ^60^ were used to compare between species.

Orthologous genes between coral species were determined using the predicted protein sequences from published reference genomes for *A. millepora* ^58^ and the *G. fascicularis* genome from this study. We identified reciprocal best BLAST hits by running BLAST (version 2.9.0, e-value threshold of 1e-6) ^36^ between predicted proteins and selecting only the gene pairs that were the best hit between the two species with custom code.

The 500 bp upstream of each gene in the genomes was identified with a script from ref. ^66^ and used as input for FIMO analysis ^67^ using default settings to identify the canonical *HSF1* binding motif (ID: MA0486 in the JASPAR database ^68^) as in ref. ^66^ (Figure 2G). Frequency of putative HSF1 binding motifs was then quantified per gene for further analysis.

**Design and synthesis of single-guide RNAs**

The *G. fascicularis* *HSF1* gene model sequence was analyzed for potential sgRNA sites by identifying all sequences in exons 2 – 8 that matched the pattern N_20_GG using the online CRISPR design tool CHOPCHOP ^69^. Sites that had matches elsewhere in the *G. fascicularis* genome were identified using Bowtie2 ^30^ and removed. As previously described, we chose two sgRNA target sites that were far enough downstream to avoid any potential alternative transcription start sites and far enough upstream that any mutations would lead to a non-functional protein product ^6,7,70^. The two sgRNAs chosen were purchased from Synthego as synthetic sgRNA (1.5 nmol) with a chemical modification (2'-O-Methyl at 3 first and last bases, 3' phosphorothioate bonds between first 3 and last 2 bases). Each lyophilized sgRNA was resuspended in 24.3 µL of nuclease-free H_2_O to a final concentration of 2 µg µL^-1^ and stored at -20°C until complexing with Cas9.

**Microinjection of sgRNA/Cas9 complexes**

Microinjection of sgRNA/Cas9 complexes was performed similarly as described by refs. ^6,7,70^, with slight modifications. Briefly, the Alt-R S.p. Cas9 Nuclease (Integrated DNA Technologies) was first diluted to 3 µg µL^-1^ with Cas9 working buffer (CWB) (4.53 mL nuclease-free H_2_O, 100 µL 1 M Hepes pH 7.5, 375 µL 2 M KCL, and 1 N NaOH to a final pH of 7.5). The two sgRNAs were complexed separately to Cas9 by mixing 1.5 µL Cas9 (3 µg µL^-1^) and 1.5 µL sgRNA (2 µg µL^-1^) and incubating at 37 °C for 15 min. sgRNA/Cas9 complexes were co-injected with a combination of both the fluorescently labeled Alex Fluor 488-dextran (Invitrogen) and 300 ng µL^-1^ mTQ2 mRNA. The 488-dextran was used for visualization during injection, while mTQ2 expression was used to identify positively injected individuals the following day. The final injection mix was as follows: 2.5 µL of sgRNA 1/Cas9 complex, 2.5 µL of sgRNA 2/Cas9 complex, 2.5 µL CWB, 0.5 µL 488-dextran, and 0.4 µL of mTQ2 mRNA (300 ng µL^-1^ final concentration). Single-cell zygotes were injected as previously described by refs. ^6,7^. For experiments using 488-dextran only (Figure S4), mTQ2 mRNA was omitted from the injection mix. Approximately 24 hpf, larvae were scored as successfully injected if fluorescence under a CFP filter was detected (Figure S3; left column). Larvae were scored as unsuccessfully injected if fluorescence under a CFP filter was absent (Figure S3; right column). Scoring was performed using a Leica M165 stereo microscope.

**Quantification of CRISPR/Cas9-based mutagenesis in *G. fascicularis***

To assess mutation frequencies for all heat stress experiments (Figures 3B-3F, S4B, and S4C), larvae of each injection class in the 27 °C treatment group were fixed in 100% ethanol and stored at -20 °C until processing after the completion of the experiments. DNA extractions for larvae were performed as previously described by ref. ^6^. Briefly, individual ethanol-fixed larvae were placed in sterile 1.5 mL centrifuge tubes. Excess ethanol was removed and 750 µL of freshly prepared lysis buffer (100 mM Tris; pH 9.0, 100 mM NaCl, 100 mM EDTA, 1% SDS) and 20 μL of Proteinase K (20 mg mL^-1^; Zymo) were added to each tube and incubated at 65 °C for 2 hours. Following incubation, 187.5 µL of 5 M potassium acetate was added to each tube, the tubes were vortexed and then incubated on ice for 10 min. Samples were centrifuged at 16,100 rcf for 20 min. Eight hundred microliters of the supernatant were removed and transferred to a new tube containing 700 µL of 100% isopropanol. Ten microliters of molecular-grade glycogen (Thermo Fisher Scientific) were added to the tube and incubated at room temperature for 30 min. DNA was then pelleted by centrifugation at 16,100 rcf for 15 min and washed with 70% ethanol. Finally, the supernatant was removed, the pellet was air dried for 5 min at ambient temperature, and the sample was resuspended in 20 µL 10 mM Tris pH 8.5. DNA samples were stored at -20 °C until processing.

To generate PCR products for Sanger sequencing, we amplified a single 781-bp region around both guide sites. Due to the proximity of the sgRNA sites, only one large amplicon was required for estimating the mutation frequency across both guide sites. Twenty-five microliter PCR reactions were set up using GoTaq® Green Master Mix (Promega) and amplicon specific primers (Table S8) as follows: 12.5 µL 2x GoTaq Master Mix, 0.5 µL forward primer (10 µM), 0.5 µL reverse primer (10 µM), 10.5 µL nuclease-free H_2_O, and 1 µL genomic DNA. PCR cycling was performed as follows: 1 cycle of 95 °C for 2 min; 34 cycles of [95 °C for 30 s, 56 °C for 45 s, 72 °C for 24 s]; one cycle of 72 °C for 5 min. The resulting PCR product was verified using gel electrophoresis and then purified using the DNA Clean & Concentrator - 5 kit (Zymo) following the manufacturer’s instructions. Finally, the cleaned PCR product (spanning both sgRNA sites) was cloned using the Topo TA Cloning Kit (Invitrogen) and 10 clones for each larva were Sanger sequenced using the M13 reverse primer (Table S8). The PCR, cloning, and sequencing of *G. fascicularis* larvae followed similar protocols to those published previously for *A. millepora* ^6^.

**Assessment of thermal tolerance in *G. fascicularis* with mutations in *HSF1***

Heat stress experiments of *G. fascicularis* larvae were performed as described in ref. ^6^. Briefly, at 3 dpf, larvae of each injection class (uninjected and those sorted as successful injected with either Cas9 only or sgRNA/Cas9 complexes) were placed into separate wells of 96-well plates containing 200 µL ASW (35 ppt) and sealed with PCR sealing film (BioRad). For the heat stress experiment comparing heat tolerance of *A. millepora* to that of *G. fascicularis* (Figures 3C-3F), the three independent replicates were from injections across three spawning days, as in ref. ^6^. For the heat stress experiment comparing injection mixes with and without mTQ2 mRNA (Figure S4), a single replicate of each injection class was injected during one spawning day. Plates were placed at 27 °C or 34 °C with 12h: 12 h light: dark cycles at 25 µmol photons m^−2^ s^−1^ and scored for survival every 24 h for 96 h. As each well contained 200 µL, the heat ramp to 34 °C was almost immediate. A larva was scored as alive if it was swimming, rotating, or had intact tissue structure. A larva was scored as dead when there was no discernable tissue structure or all tissue had disintegrated. Incubator temperatures were continuously monitored using submersible loggers in 1 L artificial seawater (HOBO pendant; Onset Computer Corp).

**Collection permits:**

CITES export permit numbers: PWS2021-AU-001696, PWS2021-AU-001696-000009, PWS2022-AU-001440, PWS2023-AU-001827

Figures

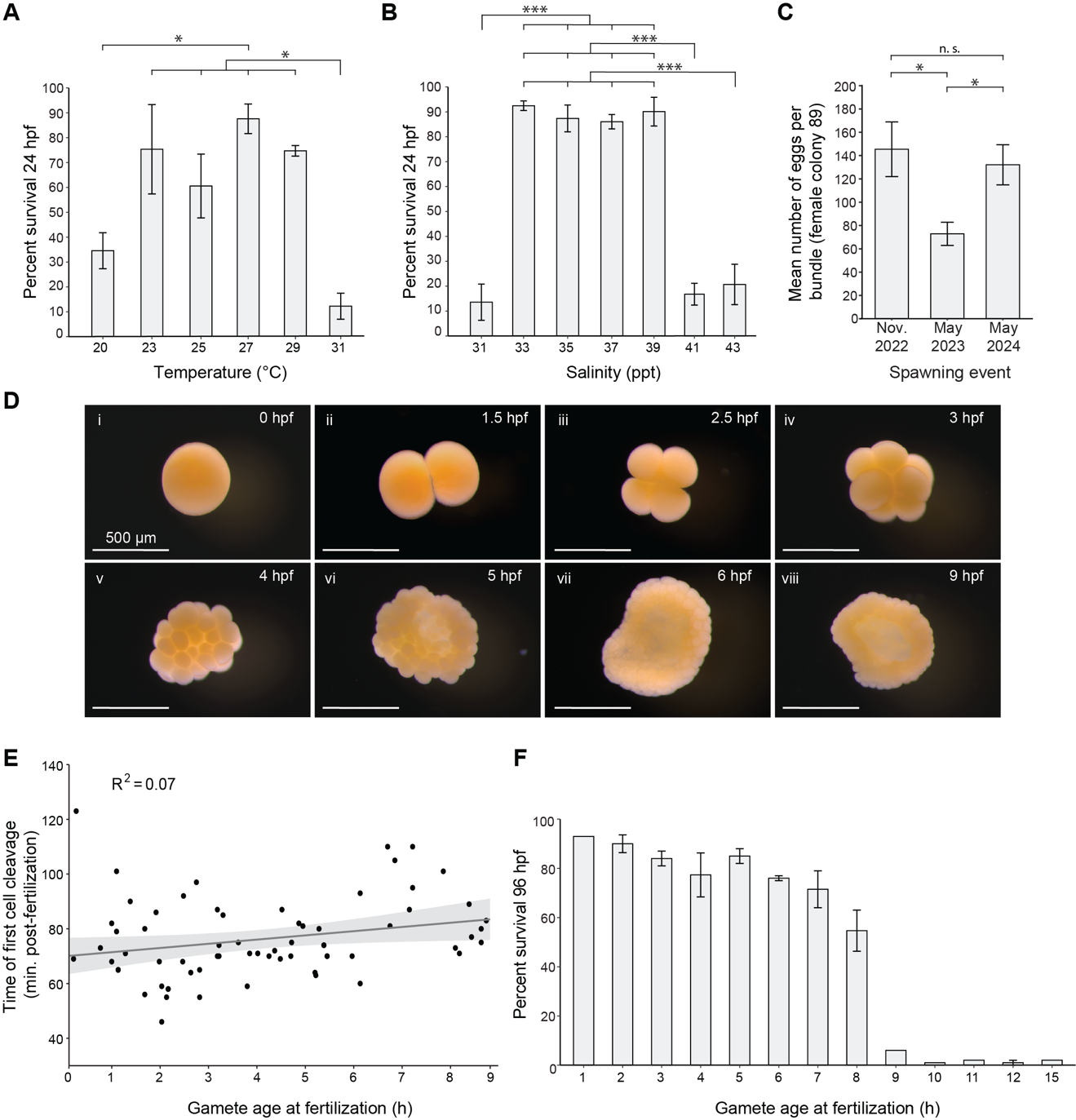

**Figure S1. Fertilization of gametes and development and survival of larvae after induced laboratory spawning. (A)** Temperature impacts on embryo survival at 24 hpf. Fertilizations were kept at each temperature until quantification. **(B)** Salinity impacts on embryo survival at 24 hpf. Fertilizations were kept at each salinity until quantification. **(C)** Mean number of eggs per bundle from a single female after a 12-month gametogenic cycle, then a reduced 6-month gametogenic cycle, before reverting back to a 12-month gametogenic cycle. **(D)** Embryonic development of *G. fascicularis* zygotes between 0 and 9 hpf. **(E)** Minutes post-fertilization of first cell cleavage of zygotes across 9 hours of staggered fertilizations. **(F)** Percent survival at 96 hpf of larvae across 15 h of staggered fertilizations. Survival at 1, 9, 10, 11, and 15 h contain a single fertilization and the remaining time points each contain two fertilization replicates. In A - C, black bars indicate standard error. In A – B, statistical significances were tested by a one-way ANOVA followed by comparisons using the Tukey HSD post hoc test (* = p ≤ 0.05; ** = p ≤ 0.001; *** = p ≤ 0.0001). In C, statistical significances were tested by a Kruskal-Wallis test followed by a Dunn’s multiple comparison test with Bonferroni correction. (* = p ≤ 0.05). In E, the grey line indicates a linear regression with standard error (grey shading).

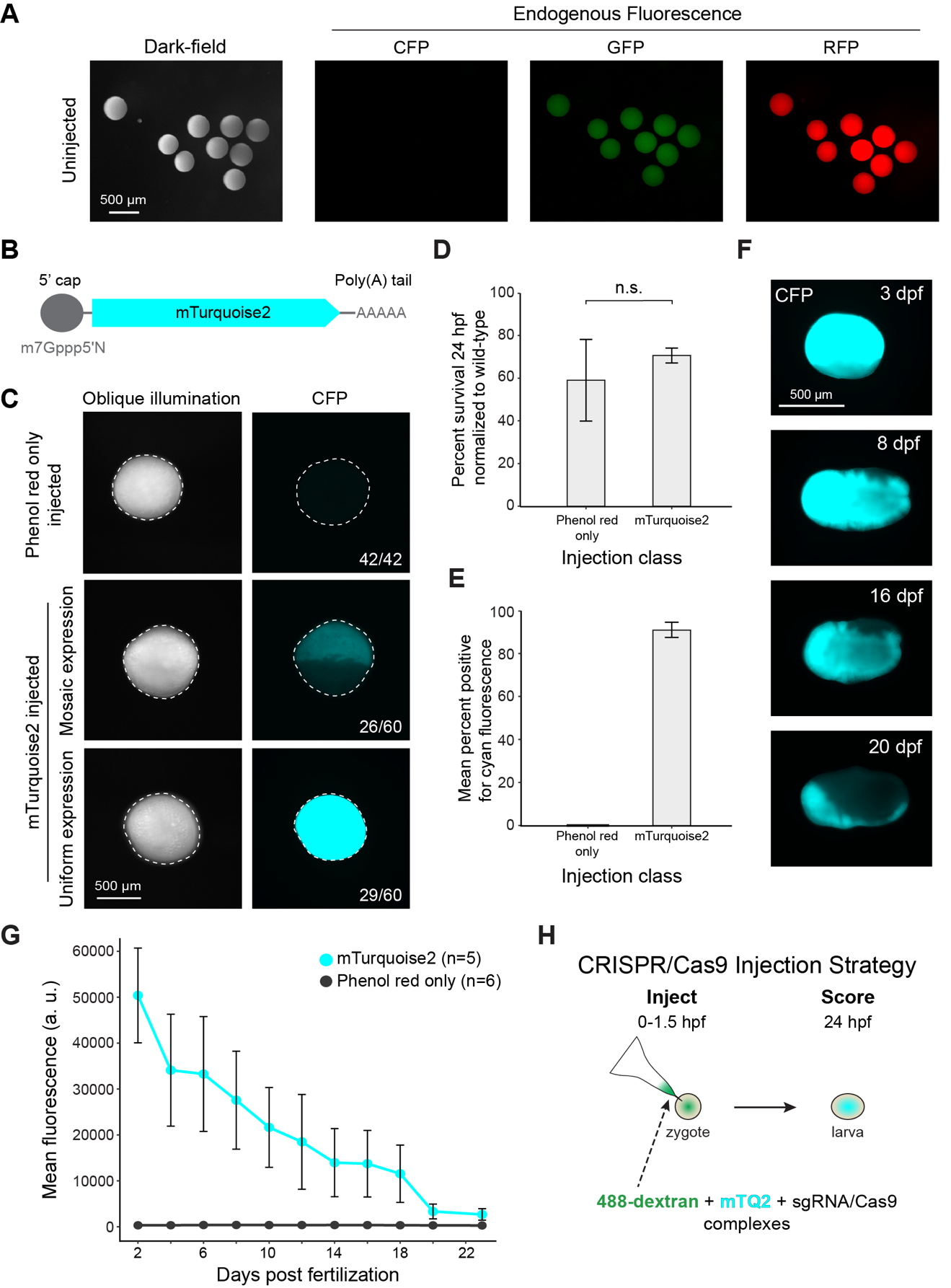

**Figure S2. mRNA overexpression and CRISPR/Cas9 injection strategy in *Galaxea fascicularis*. (A)** Endogenous fluorescence of uninjected *G. fascicularis* eggs. **(B)** Schematic representation of capped mRNA encoding a cyan fluorescent protein, mTurquoise2 (mTQ2) used for injection into 1-cell zygotes. **(C)** Representative images of larvae at 24 hpf after injection at the 1-cell stage with phenol red only (top) or with phenol red and 800 ng µL^-1^ mTQ2 mRNA (center and bottom). Numbers on the bottom right indicate the fraction of total larvae having each expression pattern. **(D)** Mean percent survival at 24 hpf of larvae injected at the 1-cell stage with phenol red only or phenol red and 800 ng µL^-1^ mTQ2 mRNA. Survival is normalized to wild-type survival at 24 hpf. Statistical significance was measured using a two sample T-test (n.s. p ≥ 0.05). **(E)** Percent of individuals injected with phenol red only or phenol red and 800 ng µL^-1^ mTQ2 mRNA from (D) scored as positively injected for cyan fluorescence approximately 24 hpf. **(F)** Fluorescence images in CFP of a larva scored as positively injected at 24 hpf over 20 days. **(G)** Mean fluorescence (arbitrary units; a. u.) over 23 days of larvae injected at the 1-cell stage with either phenol red only or with phenol red and 800 ng µL^-1^ mTQ2 mRNA. The mTQ2 larvae used in this experiment were those scored as uniformly fluorescent at 24 hpf (C; bottom). **(H)** Microinjection strategy using mTQ2 mRNA to score injection success of coral zygotes. The injection mix includes 488-dextran for visualization during injection, mTQ2 mRNA for scoring positively injected larvae approximately 24 hpf, and sgRNA/Cas9 complexes for genome editing. In D, E, and G black bars show standard error. In C and F larvae were imaged while swimming.

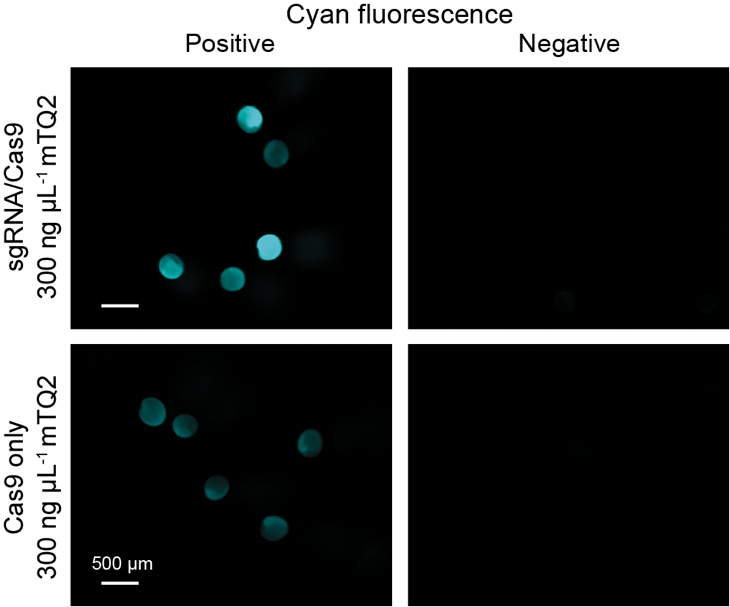

**Figure S3.**  **Scoring of injected animals at 24 hpf using cyan fluorescence.** Representative fluorescence images of *G. fascicularis* larvae at approximately 24 hpf injected at the 1-cell stage with either sgRNA/Cas9 complexes targeting *HSF1* and 300 ng µL^-1^ mTQ2 mRNA or Cas9 only and 300 ng µL^-1^ mTQ2 mRNA. Larvae were scored as successfully injected if fluorescence under a CFP filter was detected (left column). Larvae were scored as unsuccessfully injected if fluorescence under a CFP filter was absent or negligible (right column).

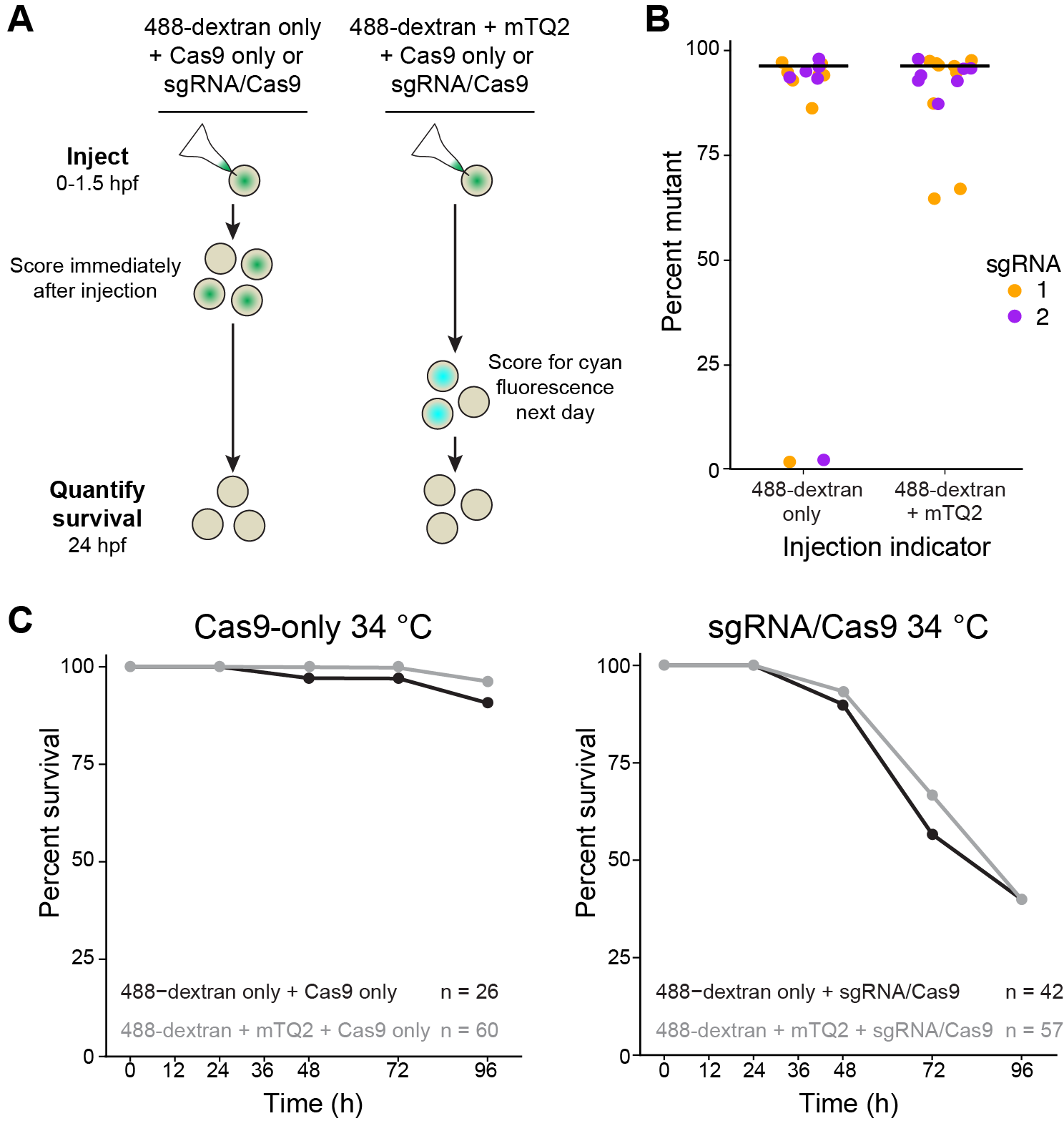

**Figure S4. Addition of mTQ2 mRNA in the injection mix does not alter larvae survival in heat when targeting *HSF1* with sgRNA/Cas9 complexes in *G. fascicularis*. (A)** 1-cell zygotes injected with mixtures of 488-dextran and Cas9 only or sgRNA/Cas9 complexes (left) were immediately scored after injection for green fluorescence above background endogenous fluorescence. Survival of these animals was quantified at 24 hpf. 1 cell-zygotes injected with mixtures of 488-dextran with 300 ng µL^-1^ mTQ2 mRNA and Cas9 only or sgRNA/Cas9 complexes (right) were scored for cyan fluorescence the following day. Survival of these animals was quantified at 24 hpf. Only a single injection replicate for each condition was performed during one spawning day. **(B)** Mutation percentages at two guide sites targeting *HSF1* of larvae from (A) scored as positively injected for each injection indicator. **(C)** Percent survival of larvae injected at the 1-cell stage with either a mixture of 488-dextran only (black) or 488-dextran and 300 ng µL^-1^ mTQ2 mRNA (grey) and Cas9 only (left) or sgRNA/Cas9 complexes targeting *HSF1* (right) during 96 h of heat stress. Experiments began when the larvae were 3 dpf. Only larvae scored as positively injected were included in the experiment.

Tables

**Table S1.** Source locations of coral colonies used in this study.

| **Species** | **Date added** | **Number of colonies received;** number distributed to aquaria | **Source reef location** | **Source location**  **(Degrees, decimal minutes)** | **Historical summer mean temperature (°C)**^(a)^ | **Historical summer maximum temperature (°C)**^(a)^ |
| --- | --- | --- | --- | --- | --- | --- |
| *Galaxea fascicularis* | Oct. 2021 | **19**; 10 to System 1,  9 to System 2 | Yamacutta | S 17° 54.347' E 146° 37.271' | 28.8 | 30.1 |
|  |  |  | Batt | S 16° 27.118' E 145° 51.153' | 28.7 | 30.5 |
|  |  |  | Tongue | S 16° 21.178' E 145° 54.996' | 29.2 | 30 |
|  |  |  | Spur | S 16° 24.208' E 146° 02.750' | 28.9 | 30.1 |
|  |  |  | Nicholas | S 16° 29.681' E 146° 06.140' | 28.9 | 30.1 |
|  |  |  | Pixie | S 16° 32.826' E 145° 51.774' | 28.9 | 30.1 |
|  |  |  | Oyster | S 16° 39.050' E 145° 56.382' | 28.6 | 30.3 |
|  |  |  | Vlasoff | S 16° 38.846' E 145° 38.846' | 28.6 | 30.3 |
|  |  |  | Arlington | S 16° 41.152' E 146° 00.247' | 28.6 | 30.3 |
|  | Feb. 2022 | **73**; 14 to System 1,  14 to System 2,  22 to System 3,  23 to System 4 | Otter | S 18° 00.035' E 146° 37.850' | 28.8 | 30.1 |
|  |  |  | Yamacutta | S 17° 52.785' E 146° 37.959' | 28.8 | 30.1 |
|  |  |  | Arlington | S 16° 39.928' E 146° 2.403' | 28.6 | 30.3 |
|  |  |  | Pretty Patches | S 16° 37.895' E 146° 3.386' | 28.6 | 30.3 |
|  |  |  | Vlassoff | S 16° 39.968' E 145° 59.873' | 28.6 | 30.3 |
|  | Sept. 2022 | **13** to System 4 | Elford | S 16° 54.841' E 146° 14.766' | 28.7 | 30.1 |
|  | Nov. 2023 | **12** to System 3 | Pixie | S 16° 32.826' E 145° 51.774' | 28.9 | 30.1 |
|  |  |  | Batt | S 16° 27.118' E 145° 51.153' | 28.7 | 30.5 |
|  |  |  | Tongue | S 16° 21.178' E 145° 54.996' | 29.2 | 30 |
|  |  |  | Spur | S 16° 24.208' E 146° 02.750' | 28.9 | 30.1 |
|  |  |  | Onyx | S 16° 26.500' E 146° 3.966' | 28.9 | 30.1 |
|  |  |  | Nicholas | S 16° 29.681' E 146° 06.140' | 28.9 | 30.1 |
|  |  |  | Pretty Patches | S 16° 37.895' E 146° 3.386' | 28.6 | 30.3 |
|  |  |  | Vlasoff | S 16° 39.968' E 145° 59.873' | 28.6 | 30.3 |
| *Acropora millepora* | -^(b)^ | - | Backnumbers | S 18° 28.925' E 147° 11.655' | 28.4 | 29.8 |
|  | -^(b)^ | - | Palm Island | S 18° 41.670' E 146° 36.323' | 28.5 | 30.1 |
|  | -^(b)^ | - | Falcon Island | S 18° 46.543' E 146° 32.1' | 28.9 | 30.4 |
| *Acropora millepora* | -^(c)^ | - | Palm Island | S 18° 41.670' E 146° 36.323' | 28.5 | 30.1 |
| ^(a)^ Temperature values are from the Australian Institute of Marine Science's Long-Term Monitoring Programs as compiled by ref. ^71^. Data were accessed through the GBR Temperature Dashboard hosted by ref. ^72^. In cases where reef data were unavailable, the next closest reef data were used. | | | | | | |
| ^(b)^ *A. millepora* colonies were collected from the Great Barrier Reef and transported to the National Sea Simulator at the Australian Institute of Marine Science in November 2018. | | | | | | |
| ^(c)^ *A. millepora* colonies were collected from the Great Barrier Reef and transported to the National Sea Simulator at the Australian Institute of Marine Science in November 2023. | | | | | | |

**Table S2.** Gametogenic length and spawning success of laboratory reared *Galaxea fascicularis* coral colonies

| **System** | **Gametogenic Length in Laboratory (months)**^(a)(b)^ | **Laboratory Spawning Event** | **Total Number of Colonies from First Spawning Cohort** | **Percent Total Colonies Spawned** | **Female Colonies Spawned**^(d)^ | **Male Colonies Spawned**^(d)^ |
| --- | --- | --- | --- | --- | --- | --- |
| System 1 | Wild | Nov. 2021 | 10 | 100.0 | 5 | 5 |
|  | 11 | Oct. 2022 | 10 | 100.0 | 5 | 5 |
|  | 12^*^ | Oct. 2023 | 10 | 80.0 | 4 | 4 |
|  | 12 | Oct. 2024 | 10 | 90.0 | 4 | 5 |
| System 2 | Wild | Nov. 2021 | 9 | 88.9 | 3 | 5 |
|  | 8 | July 2022^(c)^ | 9 | 22.2 | 0 | 2 |
|  | 9 | Aug. 2022^(c)^ | 9 | 77.8 | 3 | 4 |
|  | 12^*^ | Aug. 2023 | 7 | 100.0 | 2 | 5 |
|  | 12 | Aug. 2024 | 7 | 85.7 | 2 | 4 |
| System 3 | 14 | Jan. 2023^(c)^ | 22 | 63.6 | 5 | 8 |
|  | 15 | Feb. 2023^(c)^ | 22 | 31.8 | 1 | 6 |
|  | 12^*^ | Feb. 2024 | 18 | 66.7 | 3 | 9 |
|  | 12 | Feb. 2025 | 16 | 62.5 | 3 | 7 |
| System 4 | 12 | Nov. 2022 | 36 | 86.1 | 13 | 18 |
|  | 6 | May. 2023^(c)^ | 34 | 8.8 | 2 | 1 |
|  | 7 | Jun. 2023^(c)^ | 34 | 11.8 | 2 | 2 |
|  | 12^*^ | May. 2024 | 31 | 25.8 | 4 | 4 |
|  | 12 | May. 2025 | 27 | 48.1 | 7 | 6 |
| ^(a)^ 'Wild' indicates coral colonies that arrived in the laboratory and spawned less than one month later. | | | | | | |
| ^(b)^ Asterisks indicate the first spawning event after the cycle shifts | | | | | | |
| ^(c)^ Indicates months where a split spawning occurred | | | | | | |
| ^(d)^ Coral colony sex was determined after the first spawning event. | | | | | | |

**Table S3.** Characteristics of the Carnegie Primary *Galaxea fascicularis* genome.

| **Metric** | **Carnegie Primary** |
| --- | --- |
| Expected genome length (bp)^(a)^ | 503,331,751 |
| Heterozygosity rate (%)^(a)^ | 2.04 |
| Percent repetitive^(b)^ | 47.22 - 49.56 |
| Percent GC^(b)^ | 39.9 |
| Number of genes^(c)^ | 42,035 |
| Number of mRNA/transcript isoforms^(c)^ | 48,022 |
| Average number of isoforms per gene^(c)^ | 1.4 |
| ^(a)^ GenomeScope2 ^25,26^ |  |
| ^(b)^ RepeatMasker ^35^ |  |
| ^(c)^ Gene annotation for "Carnegie Primary" assembly (this study). | |

**Table S4.** Comparison of genome contiguity between *Galaxea fascicularis* assemblies

| **Assembly** | **Number of scaffolds** | **Assembly size (bp)** | **Maximum scaffold length (bp)** | **N50 (bp)** | **L50** | **NG50 (bp)**^(a)^ | **LG50**^(a)^ |
| --- | --- | --- | --- | --- | --- | --- | --- |
| Carnegie Primary^(b)^ | 198 | 507,902,524 | 12,998,333 | 4,809,431 | 35 | 4,809,431 | 35 |
| Gfas1.0^(c)^ | 11,269 | 334,165,880 | 874,137 | 87,933 | 1,007 | 34,693 | 2,539 |
| jaGalFasc^(d)^ | 178 | 416,866,986 | 44,590,768 | 33,853,504 | 6 | 29,797,721 | 7 |
| ^(a)^ Genome size expectation, G, was used for calculations. The genome size used was predicted by GenomeScope2 ^25,26^ based on the k-mer analysis: 503,331,751 bp. | | | | | | | |
| ^(b)^ This study. |  |  |  |  |  |  |  |
| ^(c)^ Ref. ^18,19^ |  |  |  |  |  |  |  |
| ^(d)^ Ref. ^73^ |  |  |  |  |  |  |  |

**Table S5.** Comparison of genome completeness between *Galaxea fascicularis* assemblies.

| **Assembly** | **Percent of 22,418 Gfas1.0**^(a)^ **genes successfully lifted over to assembly**^(b)^ | **Percent of 954 Metazoan BUSCOs**^(c)^ **found in genome (complete and fragmented)** | **Percent of 255 Eukaryotic BUSCOs found in genome (complete and fragmented)** | **Read mapping completeness** | **Percent of 954 Metazoan BUSCOs found in proteome (complete and fragmented)**^(d)^ | **Percent of 255 Eukaryotic BUSCOs found in proteome (complete and fragmented)**^(d)^ |
| --- | --- | --- | --- | --- | --- | --- |
| Carnegie Primary^(e)^ | 96.93 | 96.3 | 98 | 98.5 | 97.1 | 97.6 |
| Gfas1.0^(a)^ | . | 95.6 | 97.6 | 91.4 | 92.1 | 94.9 |
| jaGalFasc^(f)^ | 93.72 | 93.1 | 95.3 | 95.14 | . | . |
| ^(a)^ Ref. ^18,19^ |  |  |  |  |  |  |
| ^(b)^ Genes lifted over by Liftoff ^24^. Score not reported for Gfas1.0 because it is arbitrarily 100% as it is the source of the genes being lifted over. | | | | | |  |
| ^(c)^ BUSCO ^23^ |  |  |  |  |  |  |
| ^(d)^ jaGalFasc does not have an associated annotation. | | | |  |  |  |
| ^(e)^ This study. |  |  |  |  |  |  |
| ^(f)^ Ref. ^73^ |  |  |  |  |  |  |

**Table S6.** Consensus sequence quality values (QV) of the Carnegie Primary *Galaxea fascicularis* genome.

| **Assembly** | **Merqury QV**^(a)^ | **Merfin QV**^(b)^ | **Yak QV**^(c)^ | **Read Map QV**^(d)^ | **Median QV**^(e)^ |
| --- | --- | --- | --- | --- | --- |
| Carnegie Primary^(f)^ | 33.12 | 30.44 | 33.63 | 35.04 | 33.40 |
| To interpret, QV scores are -10*log10( error rate ). Thus, QV scores of 10, 20, 30, and 40 correspond to 1 error in 10 bp, 100 bp, 1 kb, and 10 kb, respectively. | | | | | |
| ^(a)^ Merqury ^28^ |  |  |  |  |  |
| ^(b)^ Merfin ^27^ |  |  |  |  |  |
| ^(c)^ Yak ^74^ |  |  |  |  |  |
| ^(d)^ Read map QVs were computed from the error rate of read alignments after subtracting out the error due to base calling and heterozygosity. | | | | | |
| ^(e)^ The assembly QV was estimated by four different approaches. The median QV is the median of those four QV values. | | | | | |
| ^(f)^ This study. |  |  |  |  |  |

**Table S7.** Number and survival of *Galaxea fascicularis* zygotes injected with mTurquoise2 mRNA

|  | **Phenol red dye only** | | |  | **mRNA (800 ng µL^-1^ mTQ2) and**  **phenol red dye** | | |
| --- | --- | --- | --- | --- | --- | --- | --- |
|  | **Exp. 1** | **Exp. 2** | **Exp. 3** |  | **Exp. 1** | **Exp. 2** | **Exp. 3** |
| Number of zygotes for which injection was attempted | 63 | 75 | 81 |  | 75 | 91 | 75 |
| Percent of individuals surviving 24 h post-fertilization normalized to wild-type | 22.7 | 87.5 | 67.0 |  | 67.0 | 74.5 | 73.7 |
| Percent of surviving individuals with mosaic or uniform cyan fluorescence | 0 | 0 | 0 |  | 95 | 88.9 | 90.9 |

**Table S8.** Oligonucleotides used in this study.

| **Oligonucleotide name** | **Purpose** | **Oligonucleotide sequence (5' to 3')** |
| --- | --- | --- |
| Pcmv_end_seq_fwd | Amplification from Turquoise2 plasmid sequence to make mRNA | GCAAATGGGCGGTAGGCG |
| M13 reverse^(a)^ |  | CAGGAAACAGCTATGAC |
| sgRNA 1 | sgRNAs used in *G. fascicularis* ^(b)^ | GAGATCTGAGAAAGATGAGT |
| sgRNA 2 |  | AATGACGTCCAGGATATGAA |
| Primer 1 (NS355) | Amplification of genomic DNA fragments for sequencing ^(c)^ | GCAGTCCAGGAATAGAATTGCAC |
| Primer 2 (AT146) |  | TCCACCAGTTCATTCCACAA |
| ^(a)^ M13 reverse was also used to Sanger sequence *G. fascicularis* larvae | | |
| ^(b)^ sgRNA guides were from Synthego (chemical modifications: 2'-O-Methyl at 3 first and last bases, 3' phosphorothioate bonds between first 3 and last 2 bases) | | |
| ^(c)^ A single 781-bp amplicon product across both sgRNA guide sites used for sequencing | | |

**Table S9.** Number and survival of *Galaxea fascicularis* zygotes injected with Cas9 only and sgRNA/Cas9.

|  | **Cas9 only (300 ng µL^-1^ mTQ2 mRNA)** | | |  | **sgRNA/Cas9 (300 ng µL^-1^ mTQ2 mRNA)** | | |
| --- | --- | --- | --- | --- | --- | --- | --- |
|  | **Exp. 1** | **Exp. 2** | **Exp. 3** |  | **Exp. 1** | **Exp. 2** | **Exp. 3** |
| Number of zygotes for which injection was attempted | 220 | 177 | 408 |  | 205 | 173 | 353 |
| Percent of individuals surviving ~24 h post-fertilization normalized to wild-type | 62.8 | 55.2 | 30.0 |  | 73.2 | 38.5 | 93.5 |
| Percent of surviving individuals scored as positive for cyan fluorescence | 84.7 | 67.7 | 86.4 |  | 70.7 | 66.7 | 89.7 |

**Table S10.** Mutations in *G. fascicularis* larvae injected with sgRNA/Cas9 complexes targeting *HSF1*.

|  |  |  | **Sequenced Clones**^(c)^ | |  |  |  |  |  | |  | |  | |  | |
| --- | --- | --- | --- | --- | --- | --- | --- | --- | --- | --- | --- | --- | --- | --- | --- | --- |
| **Experiment**^(a)^ | **Animal**^(b)^ | **Exon** | **1** | **2** | **3** | **4** | **5** | **6** | | **7** | | **8** | | **9** | **10** | **Percent of sequences with large deletions** |
| 1 | 1 | 3 | -397, +1 | complex | -397, +1 | -397, +1 | -397, +1 | complex | | -397, +1 | | -397, +1 | | -397, +1 | -397, +1 | 100 |
|  |  | 4 | -397, +1 | complex | -397, +1 | -397, +1 | -397, +1 | complex | | -397, +1 | | -397, +1 | | -397, +1 | -397, +1 |  |
|  | 2 | 3 | -399, +8 | complex | -399, +8 | complex | -399, +8 | -399, +8 | | complex | | -399, +8 | | -399, +8 | -399, +8 | 100 |
|  |  | 4 | -399, +8 | complex | -399, +8 | complex | -399, +8 | -399, +8 | | complex | | -399, +8 | | -399, +8 | -399, +8 |  |
|  | 3 | 3 | -2 | -407 | -407 | -407 | -402 | -398 | | -2 | | -398 | | -4 | -2 | 60 |
|  |  | 4 | -9 | -407 | -407 | -407 | -402 | -398 | | -9 | | -398 | | -9 | -9 |  |
|  | 4 | 3 | -28 | -28 | -28 | -28 | -28 | -28 | | -27 | | -28 | | -28 | -28 | 0 |
|  |  | 4 | -26 | -26 | -26 | -26 | -26 | -26 | | -4 | | -26 | | -26 | -26 |  |
|  | 5 | 3 | -406 | -406 | -423 | -406 | -423 | -406 | | -406 | | -406 | | -406 | -423 | 100 |
|  |  | 4 | -406 | -406 | -423 | -406 | -423 | -406 | | -406 | | -406 | | -406 | -423 |  |
|  | 6 | 3 | -409 | -409 | -7 | -409 | -409 | -7 | | -37 | | -409 | | -409 | -7 | 60 |
|  |  | 4 | -409 | -409 | -6 | -409 | -409 | -6 | | -9 | | -409 | | -409 | -7 |  |
|  | 7 | 3 | +6 | -4 | -4 | -4 | +6 | complex | | complex | | -4 | | -4 | complex | 0 |
|  |  | 4 | -5 | -5 | -5 | 1 SNP | -5 | complex | | complex | | +6 | | -5 | complex |  |
|  | 8 | 3 | -4 | -4 | -4 | -4 | -4 | -4 | | +20 | | +20 | | +6 | complex | 0 |
|  |  | 4 | -9 | -9 | -9 | -9 | -9 | -9 | | complex | | complex | | -5 | -5 |  |
|  | 9 | 3 | -409, +6 | -409, +6 | -409, +6 | -409, +6 | -409, +6 | -409, +6 | | -409, +6 | | -409, +6 | | -409, +6 | -409, +6 | 100 |
|  |  | 4 | -409, +6 | -409, +6 | -409, +6 | -409, +6 | -409, +6 | -409, +6 | | -409, +6 | | -409, +6 | | -409, +6 | -409, +6 |  |
|  | 10 | 3 | -32 | -5 | -22 | -22 | -5 | -32 | | -22 | | -32 | | -5 | -22 | 0 |
|  |  | 4 | -2 | -5 | -5 | -5 | -5 | -2 | | -5 | | -2 | | -5 | -5 |  |
|  | 11 | 3 | -9 | -9 | -17 | -9 | -401, +5 | complex | | -4 | | -401, +5 | | -4 | -9 | 20 |
|  |  | 4 | -14 | -14 | -14 | -14 | -401, +5 | -5 | | -4 | | -401, +5 | | -4 | -13 |  |
|  | 12 | 3 | -414, +7 | WT | WT | WT | WT | -4 | | WT | | WT | | +2 | WT | 10 |
|  |  | 4 | -414, +7 | WT | WT | WT | WT | WT | | -6 | | -9 | | WT | WT |  |
|  | 13 | 3 | -397 | WT | 1 SNP | -419, +73 | WT | WT | | -20 | | WT | | -419, +73 | 1 SNP | 30 |
|  |  | 4 | -397 | WT | -9 | -419, +73 | -10 | +2, -20 | | WT | | +2, -20 | | -419, +73 | WT |  |
|  | 14 | 3 | -405, +8 | -2 | -2 | -405, +8 | -405, +8 | -405, +8 | | -405, +8 | | -405, +8 | | -405, +8 | -405, +8 | 80 |
|  |  | 4 | -405, +8 | -4 | complex | -405, +8 | -405, +8 | -405, +8 | | -405, +8 | | -405, +8 | | -405, +8 | -405, +8 |  |
|  | 15 | 3 | WT | WT | 2 SNP | WT | WT | WT | | WT | | WT | | WT | WT | 0 |
|  |  | 4 | WT | WT | WT | complex | WT | WT | | WT | | WT | | WT | WT |  |
|  | 16 | 3 | -402, +25 | -400, +39 | complex | -18 | complex | complex | | -400, +39 | | complex | | complex | -396, +138 | 40 |
|  |  | 4 | -402, +25 | -400, +39 | complex | -9 | complex | complex | | -400, +39 | | complex | | complex | -396, +138 |  |
|  | 17 | 3 | -408, +23 | -408, +23 | -408, +23 | -408, +23 | -4 | -9 | | -408, +23 | | -4 | | -233, +4 | -503 | 60 |
|  |  | 4 | -408, +23 | -408, +23 | -408, +23 | -408, +23 | -4 | -5 | | -408, +23 | | -19, +4 | | -18 | -503 |  |
|  | 18 | 3 | -2 | -13 | complex | complex | -2 | -401, +4 | | -2 | | -4 | | complex | -401, +4 | 20 |
|  |  | 4 | complex | complex | complex | -18 | -10 | -401, +4 | | -5 | | -35 | | complex | -401, +4 |  |
|  | 19 | 3 | -322 | -44 | -322 | -322 | -322 | -322 | | -322 | | -322 | | -322 | -322 | 0 |
|  |  | 4 | -1, +10 | -199, +2 | -1, +10 | -1, +10 | -1, +10 | -1, +10 | | -1, +10 | | -1, +10 | | -1, +10 | -1, +10 |  |
| 2 | 1 | 3 | -406 | -40, +1 | -5 | -406 | -31 | complex | | complex | | -40, +1 | | -406 | -5 | 30 |
|  |  | 4 | -406 | -9 | -202 | -406 | -202 | -202 | | complex | | -9 | | -406 | -202 |  |
|  | 2 | 3 | -411 | -397 | -397 | -397 | -397 | -411 | | -411 | | -411 | | -411 | -411 | 100 |
|  |  | 4 | -411 | -397 | -397 | -397 | -397 | -411 | | -411 | | -411 | | -411 | -411 |  |
|  | 3 | 3 | -49 | -427, +25 | -427, +25 | complex | -427, +25 | -427, +25 | | -427, +25 | | -427, +25 | | -427, +25 | -427, +25 | 80 |
|  |  | 4 | -5 | -427, +25 | -427, +25 | -8 | -427, +25 | -427, +25 | | -427, +25 | | -427, +25 | | -427, +25 | -427, +25 |  |
|  | 4 | 3 | -5 | -10 | -9 | -9,+1 | +4,-40 | -9 | | -5 | | -9 | | -9 | +4,-40 | 0 |
|  |  | 4 | -5 | -5 | -5 | -5 | -5 | -5 | | -5 | | -62 | | -62 | -21 |  |
|  | 5 | 3 | complex | complex | complex | complex | complex | -469, +27 | | complex | | complex | | -6 | complex | 80 |
|  |  | 4 | complex | complex | complex | -14 | complex | -469, +27 | | complex | | complex | | -4 | complex |  |
|  | 6 | 3 | complex | complex | +7 | complex | complex | complex | | -12 | | complex | | -12 | complex | 20 |
|  |  | 4 | -23 | complex | -23 | -18 | complex | -18 | | -18 | | -23 | | -18 | -23 |  |
|  | 7 | 3 | -402 | -402 | -402 | -402 | -402 | -8 | | -402 | | -5 | | -8 | -9, +1 | 60 |
|  |  | 4 | -402 | -402 | -402 | -402 | -402 | -14, +5 | | -402 | | -32 | | -14, +5 | -5 |  |
|  | 8 | 3 | -490 | -411 | -411 | -411 | -399, +80 | -399, +80 | | -411 | | -399, +80 | | -411 | -411 | 100 |
|  |  | 4 | -490 | -411 | -411 | -411 | -399, +80 | -399, +80 | | -411 | | -399, +80 | | -411 | -411 |  |
|  | 9 | 3 | -4 | -4 | -9 | -4 | -9 | complex | | -27, +1 | | -9 | | -27, +1 | -9 | 0 |
|  |  | 4 | -2 | -2 | complex | -2 | complex | -2 | | -4 | | -6 | | -8 | -3 |  |
| 3 | 1 | 3 | -402 | -402 | -402 | -2 | complex | -426 | | -426 | | -426 | | -426 | -426 | 80 |
|  |  | 4 | -402 | -402 | -402 | -33 | -4 | -426 | | -426 | | -426 | | -426 | -426 |  |
|  | 2 | 3 | -207 | -5 | -207 | -5 | -207 | -5 | | -13 | | -2, +1 | | -5 | -207 | 0 |
|  |  | 4 | -5 | -14 | -5 | -14 | -9 | -6 | | -14 | | -14 | | -6 | -9 |  |
|  | 3 | 3 | -6 | -2 | -2 | -2 | -2 | -2 | | -2 | | -398 | | -9 | -2 | 10 |
|  |  | 4 | complex | complex | complex | complex | complex | complex | | complex | | -398 | | -18 | complex |  |
|  | 4 | 3 | -401 | -3 | -3 | -3 | -405 | -415 | | -3 | | -405 | | -4 | -405 | 50 |
|  |  | 4 | -401 | -359 | -359 | -359 | -405 | -415 | | -359 | | -405 | | -6 | -405 |  |
|  | 5 | 3 | -27 | -21 | -2 | +10 | +10 | -14 | | -9 | | -13 | | +10 | -407 | 10 |
|  |  | 4 | -5 | -27, +2 | -71, +5 | -71, +5 | -71, +5 | +4 | | -152 | | -71, +5 | | -71, +5 | -407 |  |
|  | 6 | 3 | -2 | -68, +52 | -27 | -28 | -27 | -27 | | -4 | | -27 | | -18, +1 | -397 | 10 |
|  |  | 4 | -71, +5 | -4 | -9 | -4 | -9 | -9 | | -71, +5 | | -9 | | -71, +5 | -397 |  |
|  | 7 | 3 | -28, +1 | -4 | -542 | -22 | -413 | -15 | | -492 | | -402 | | -414, +1 | -28, +1 | 50 |
|  |  | 4 | -18 | +17 | -542 | -15 | -413 | -5 | | -492 | | -402 | | -414, +1 | -18 |  |
|  | 8 | 3 | -27 | -28 | -4 | -412 | -25 | -16 | | -9 | | -402 | | -413 | -402 | 40 |
|  |  | 4 | -14 | complex | complex | -412 | complex | complex | | -9, +1 | | -402 | | -413 | -402 |  |
|  | 9 | 3 | -2 | -28 | complex | 1 SNP | -2 | complex | | -2 | | -1, +1 | | -402 | -439 | 20 |
|  |  | 4 | complex | -4 | -4 | -6 | complex | -5 | | complex | | -37 | | -402 | -439 |  |
|  | 10 | 3 | WT | complex | -4 | -5 | -396, +33 | -27, +3 | | -405 | | -9 | | -396, +33 | -6 | 30 |
|  |  | 4 | -5 | -3 | -3 | 1 SNP | -396, +33 | -4 | | -405 | | WT | | -396, +33 | -3 |  |
|  | 11 | 3 | -407 | -13 | complex | WT | -1 | -14, +1 | | -408 | | +3 | | -3 | WT | 20 |
|  |  | 4 | -407 | -9 | -9 | -8 | complex | complex | | -408 | | -8 | | -3 | -8 |  |
| ^(a)^ Experiment numbers correspond to three independent days of *G. fascicularis* spawning and microinjection. | | | | | | | | | | | | | | | | |
| ^(b)^ Black numbers correspond to animals with uniform mTQ2 expression (Figure S2C; bottom). Red numbers correspond to animals with partial/mosaic mTQ2 expression (Figure S2C; center) | | | | | | | | | | | | | | | | |
| ^(c)^ Blue cells indicate large deletions spanning from sgRNA 1 to sgRNA 2. | | | | | | | | | | | | | | | | |

**Table S11.** Number and survival of *Galaxea fascicularis* zygotes injected with Cas9 only and sgRNA/Cas9.

|  | **488-dextran only** | |  | **488-dextran + 300 ng µL^-1^ mTQ2 mRNA** | |
| --- | --- | --- | --- | --- | --- |
|  | **Cas9 only** | **sgRNA/Cas9** |  | **Cas9 only** | **sgRNA/Cas9** |
| Number of zygotes for which injection was attempted | 67 | 183 |  | 316 | 178 |
| Percent of individuals surviving ~24 h post-fertilization normalized to wild-type | 71.6 | 57.0 |  | 80.5 | 62.0 |
| Percent of surviving individuals scored as positive for cyan fluorescence | - | - |  | 76.6 | 94.9 |

**Table S12.** Mutations in larvae injected with sgRNA/Cas9 complexes.

|  |  |  | **Sequenced Clones** | |  |  |  |  |  |  |  |  |  |
| --- | --- | --- | --- | --- | --- | --- | --- | --- | --- | --- | --- | --- | --- |
| **Injection Indicator** | **Animal** | **Exon** | **1** | **2** | **3** | **4** | **5** | **6** | **7** | **8** | **9** | **10** | **Percent of sequences with large deletions** |
| mTQ2 + | 1 | 3 | -4 | -4 | -4 | -4 | -4 | -4 | -4 | -4 | -4 | -4 | 0 |
| 488-dextran |  | 4 | -6 | -6 | -18 | -18 | -6 | -6 | -6 | -18 | -6 | -6 |  |
|  | 2 | 3 | -398 | complex | complex | +3 | -398 | -11 | -398 | -398 | -398 | -4 | 60 |
|  |  | 4 | -398 | -8 | complex | -8 | -398 | complex | -398 | -398 | -398 | -9 |  |
|  | 3 | 3 | -408, +8 | -402 | -4 | complex | complex | -408, +8 | -429 | -408, +8 | complex | -406 | 60 |
|  |  | 4 | -408, +8 | -402 | -18 | complex | -15 | -408, +8 | -429 | -408, +8 | complex | -406 |  |
|  | 4 | 3 | WT | -444 | -397, +17 | -444 | -396, +4 | -4 | WT | complex | -402 | WT | 50 |
|  |  | 4 | -8 | -444 | -397, +17 | -444 | -396, +4 | -8 | -4 | -10 | -402 | -8 |  |
|  | 5 | 3 | complex | -2 | +6 | -4 | complex | complex | -5 | -2 | -2 | -2 | 0 |
|  |  | 4 | -11 | -5 | -14 | -18 | -18 | -5 | -18 | -4 | -4 | -4 |  |
|  | 6 | 3 | +5 | -397, +5 | -244 | -405 | WT | WT | -6 | WT | complex | -397 | 30 |
|  |  | 4 | WT | -397, +5 | -23, +2 | -405 | 3 SNP | -15 | -9 | -5 | -5 | -397 |  |
|  | 7 | 3 | +2 | -411 | -411 | -403, +58 | -5 | -411 | -411 | -403, +58 | -4 | -4 | 60 |
|  |  | 4 | -14 | -411 | -411 | -403, +58 | complex | -411 | -411 | -403, +58 | -14 | -14 |  |
|  | 8 | 3 | -26 | complex | complex | -8, +1 | -26 | +62 | -4 | -2 | complex | complex | 0 |
|  |  | 4 | -15 | complex | -3 | -4 | -15 | -145 | -4 | -4 | complex | -4 |  |
|  | 9 | 3 | +12 | -402 | -279, +2 | -402 | -402 | -402 | +12 | -2 | -402 | -402 | 60 |
|  |  | 4 | -1 | -402 | -9 | -402 | -402 | -402 | -1 | -126, +1 | -402 | -402 |  |
|  | 10 | 3 | -1 | -5 | -1 | complex | WT | complex | +27 | +3 | complex | -5, +3 | 0 |
|  |  | 4 | -1, +1 | -11, +3 | -9 | complex | -6 | -5 | complex | -6 | -18 | -6 |  |
| 488-dextran | 1 | 3 | -29, +1 | -4 | -401, +21 | -404 | -400, +3 | -399, +4 | -5 | -397 | WT | -5 | 50 |
| only |  | 4 | -11 | -8 | -401, +21 | -404 | -400, +3 | -399, +4 | -3, +19 | -397 | -10 | -5 |  |
|  | 2 | 3 | -405 | complex | -4 | -405 | complex | -405 | -405 | complex | +10 | -405 | 50 |
|  |  | 4 | -405 | complex | -18 | -405 | complex | -405 | -405 | complex | complex | -405 |  |
|  | 3 | 3 | -398 | -10, +3 | -5 | -23 | -5 | -9 | -9 | complex | -5 | -4 | 20 |
|  |  | 4 | -398 | complex | complex | +1 | complex | -18 | complex | complex | complex | complex |  |
|  | 4 | 3 | -406 | complex | -398, +1 | -4 | -402 | -396 | -415 | -2 | -397 | -2 | 60 |
|  |  | 4 | -406 | complex | -398, +1 | -4 | -402 | -396 | -415 | -4 | -397 | complex |  |
|  | 5 | 3 | -34 | -397 | -4 | -25 | +6 | -301 | -4 | -301 | -5 | -397 | 20 |
|  |  | 4 | -10 | -397 | complex | complex | -9 | -44 | complex | -44 | -6 | -397 |  |
|  | 6 | 3 | -402 | -397, +37 | -9 | -397, +37 | -397, +37 | -397, +37 | -9 | -397, +37 | -397, +37 | -397, +37 | 80 |
|  |  | 4 | -402 | -397, +37 | -45 | -397, +37 | -397, +37 | -397, +37 | -26, +1 | -397, +37 | -397, +37 | -397, +37 |  |
|  | 7 | 3 | -4 | -4 | -4 | -4 | -4 | -4 | -139, +7 | -4 | -4 | -4 | 0 |
|  |  | 4 | -358 | -358 | complex | -358 | complex | -358 | -15 | -358 | -358 | -358 |  |
|  | 8 | 3 | WT | WT | WT | WT | WT | WT | WT | WT | WT | WT | 0 |
|  |  | 4 | WT | WT | WT | WT | WT | WT | WT | WT | WT | WT |  |
|  | 9 | 3 | -5 | -402 | -498 | -412 | -412 | -402 | -412 | -412 | -411 | -402 | 90 |
|  |  | 4 | -4 | -402 | -498 | -412 | -412 | -402 | -412 | -412 | -411 | -402 |  |
|  | 10 | 3 | -402 | -411 | -3 | -397 | -397 | -397 | -7 | -5 | -397 | -397 | 70 |
|  |  | 4 | -402 | -411 | -8 | -397 | -397 | -397 | complex | -5 | -397 | -397 |  |
| ^a^ Blue cells indicate large deletions spanning from sgRNA 1 to sgRNA 2. | | | | | | | | | | | | | |

72.	Environmental research, maps and data for tropical Australia | eAtlas https://eatlas.org.au/.

73.	Galaxea fascicularis genome assembly jaGalFasc40.1 NCBI. https://www.ncbi.nlm.nih.gov/datasets/genome/GCA_948470475.1/.

74.	Jiang, H., Chai, Z.-X., Chen, X.-Y., Zhang, C.-F., Zhu, Y., Ji, Q.-M., and Xin, J.-W. (2024). Yak genome database: a multi-omics analysis platform. BMC Genomics *25*, 346. https://doi.org/10.1186/s12864-024-10274-6.
